# Sphingosine kinases promote Ebola virus infection and can be targeted to inhibit filoviruses, coronaviruses, and arenaviruses using late endocytic trafficking to enter cells

**DOI:** 10.1101/2022.08.12.503750

**Authors:** Corina M. Stewart, Yuxia Bo, Kathy Fu, Mable Chan, Robert Kozak, Kim Yang-Ping Apperley, Geneviève Laroche, André Beauchemin, Gary Kobinger, Darwyn Kobasa, Marceline Côté

## Abstract

Entry of enveloped viruses in host cells requires the fusion of the viral and host cell membranes, a process that is facilitated by viral fusion proteins protruding from the viral envelope. For fusion, viral fusion proteins need to be triggered by host factors and for some viruses, such as Ebola virus (EBOV) and Lassa fever virus, this event occurs inside endosomes and/or lysosomes. Consequently, these ‘late-penetrating viruses’ must be internalized and delivered to entry-conducive intracellular vesicles. Because endocytosis and vesicular trafficking are tightly regulated cellular processes, late penetrating viruses also depend on specific host factors, such as signaling molecules, for efficient viral delivery to the site of fusion, suggesting that these could be targeted for antiviral therapy. In this study, we investigated a role for sphingosine kinases (SKs) in viral entry and found that chemical inhibition of sphingosine kinase 1 (SK1) and/or SK2 and knockdown of SK1 or SK2, inhibited entry of EBOV into host cells. Mechanistically, inhibition of SK1 and/or SK2 prevented EBOV from reaching late-endosomes and lysosomes that are positive for the EBOV receptor, Niemann Pick C1 (NPC1). Furthermore, we present evidence that suggests the trafficking defect caused by SK1/2 inhibition occurs independently of S1P signaling through cell-surface S1PRs. Lastly, we found that chemical inhibition of SKs prevents entry of other late-penetrating viruses, including arenaviruses and coronaviruses, in addition to inhibiting infection by replication competent EBOV and SARS-CoV-2 in Huh7.5 cells. In sum, our results highlight an important role played by SKs in endocytic trafficking which can be targeted to inhibit entry of late-penetrating viruses. SK inhibitors could serve as a starting point for the development of broad-spectrum antiviral therapeutics.

## Introduction

To deliver the viral genome into host cells, enveloped viruses require the fusion between the viral and cellular membranes. This process is mediated by viral fusion proteins that protrude from the viral envelope and undergo a series of conformational changes triggered by interactions with host factors and/or environmental cues such as acidic pH. For Ebola virus (EBOV), the causative agent of outbreaks of severe hemorrhagic fevers in humans, its fusion protein, the viral glycoprotein (GP), requires cleavage by low pH-dependent cathepsin proteases and interaction with the late endosomal/lysosomal Niemann-Pick C1 protein (NPC1) (1-5). The strict late endosomal/lysosomal localization of the EBOV GP triggering factors results in a need for a multistep viral entry process involving attachment, internalization, and endosomal trafficking. Attachment of EBOV to the host cell is mediated by interactions of GP to cell surface carbohydrate binding proteins such as C-type lectins, and/or by binding of phosphatidylserine (PS) on the viral envelope to cell surface PS receptors such as Tim-1 or Axl (6-8). Once attached to the cell surface, internalization via macropinocytosis or phagocytosis is triggered and is followed by endosomal trafficking to late endosomes and lysosomes where viral fusion occurs (2, 3, 9-14).

Many host proteins have been identified to be important for endosomal trafficking of EBOV, such as the small GTPase Rab7, the homotypic fusion and protein sorting (HOPS) complex, the UV Radiation Resistance Associated (UVRAG) protein, and the PIKfyve-ArPIKfyve-Sac3 trafficking complex (10, 15, 16). In addition, studies have shown that EBOV particles can stimulate signaling cascades such as the Akt/PI3K pathway, the latter being required for EBOV trafficking to NPC1 (16-19). In a previous study, we screened a kinase inhibitor library to identify other signaling pathways important for EBOV entry (18). We identified 35 compounds that inhibited EBOV entry, including inhibitors of receptor tyrosine kinases (RTKs), AMP-activated protein kinase (AMPK), and Akt, some of which have previously been shown to be important for virus internalization and trafficking (17-20). Additionally, inhibitors of sphingosine kinase 1 (SK1) and sphingosine-1-phosphate G-protein coupled receptors (S1PR) were also identified, suggesting a role for the SK-S1PR axis in EBOV GP-mediated entry (18). Interestingly, Imre and colleagues reported that a sphingosine kinase 1 activator or the overexpression of SK1 reduced EBOV-GP mediated entry, further suggesting a role for SK1 in EBOV entry, although the mechanism remains to be elucidated (21).

Sphingosine kinases 1 and 2 (SK1/2) catalyze the phosphorylation of sphingosine to sphingosine-1-phosphate (S1P). Despite their polypeptide sequence similarity, SK1 and 2 appear to have distinct physiological functions; generally, SK1 promotes cell survival and proliferation while SK2 is pro-apoptotic (22). S1P has both extracellular and intracellular targets. Extracellular S1P activates signaling pathways through a family of five S1P GPCRs (S1PR_1-5_), while intracellular S1P has been shown to modulate the activity of multiple enzymes including p-21 activated kinase 1 (Pak1), which is an important regulator of macropinocytosis (23-25). In addition, recent studies have also demonstrated a link between SK1, levels of sphingosine, and regulation of endocytic membrane trafficking (26, 27). Their putative roles in macropinocytosis and trafficking indicate that SK1 and SK2 may be entry host factors for EBOV and potential targets for antiviral therapy.

Viruses, such as EBOV, that enter cells via late endosomes or lysosomes are termed late penetrating viruses. Other examples of such viruses are Lassa fever virus (LFV) and severe acute respiratory syndrome coronaviruses -1 and -2 (SARS-CoV-1 and SARS-CoV-2). LFV entry requires low pH and interaction with the lysosomal-associated membrane protein 1 (LAMP-1) (28-31). Therefore, similarly to EBOV, LFV must be internalized and undergo endo-lysosomal trafficking to reach entry-conducive intracellular vesicles (32). As for SARS-CoV-1 and -2, the cellular localization of fusion depends on the expression and activity of cell surface proteases that are required for priming and/or triggering (33-36). More specifically, SARS-CoV-1 can utilize surface serine proteases, while SARS-CoV-2 is more promiscuous in its protease usage, having been shown to utilize both surface serine proteases and metalloproteases for entry at the cell surface (33-37). In the absence of surface or extracellular proteases, the entry of SARS CoV-1 and -2 depends on internalization and trafficking to late endosomes containing cathepsin L (36, 38-40). Although the triggering factors are different for each of these viruses, one commonality is a need for endosomal trafficking in host cells, raising the possibility that they may share the use of host trafficking factors (41).

Here, we investigated the role of sphingosine kinases in cellular entry of EBOV and other late penetrating viruses. We found that treatment with sphingosine kinase inhibitors and knockdown of sphingosine kinase 1 and 2 reduced EBOV growth and/or EBOV GP-mediated entry. More specifically, the sphingosine kinase inhibitors blocked trafficking of EBOV viral-like particles (VLPs) to NPC1 independently of S1P signaling through S1PRs. Lastly, we found that treatment with the SK inhibitors reduced entry of other late penetrating viruses and blocked cathepsin-dependent infection of SARS CoV-2. Our results suggest that SKs play a critical role in regulating endosomal trafficking and that inhibition of SK activity results in trafficking defects that prevent late-penetrating viruses from reaching entry-conducive compartments.

## Results

### Sphingosine kinase inhibitors reduce entry of EBOV VLPs

Previously, we screened a library of kinase inhibitors using murine leukemia virus (MLV) pseudotypes to identify signaling pathways required for EBOV GP-mediated entry (18). Among the hits were two compounds, PF-543 and FTY-720 (fingolimod, 2-amino-2-(2-(4-octylphenyl)-ethyl)-1,3-propanediol), that interfere with the SK/S1PR signaling axis, suggesting that this pathway plays an important role in EBOV entry. PF-543 is a SK1 and SK2 competitive inhibitor, while FTY-720 (fingolimod, 2-amino-2-(2-(4-octylphenyl)-ethyl)-1,3-propanediol) is a structural analog of sphingosine that becomes phosphorylated by SK1 and SK2, albeit more efficiently by SK2, and thus can inhibit the SKs (42, 43). We therefore sought to investigate the possible role of the SK/S1PR signaling pathway in EBOV entry by validating and characterizing the antiviral activity of these inhibitors and that of SK1-I, which is a sphingosine analogue that is highly specific to SK1 over SK2 and was not present in the screening library (44). To confirm the antiviral activity of PF-543, FTY-720, and SK1-I, we used filoviral-like particles (VLPs) generated by the co-expression of the EBOV nucleoprotein (NP), matrix protein (VP40) fused to a β-lactamase (βlam) reporter, and the glycoprotein of interest (45). We found that treatment with all three inhibitors reduced entry of VLPs harbouring EBOV GP (Figure 1A-C). Importantly, the inhibitors had no effect on entry of VLPs harbouring the vesicular stomatitis virus (VSV) G protein, indicating that the compounds target an EBOV GP-mediated entry step (Figure 1A-C).

**Figure 1.**
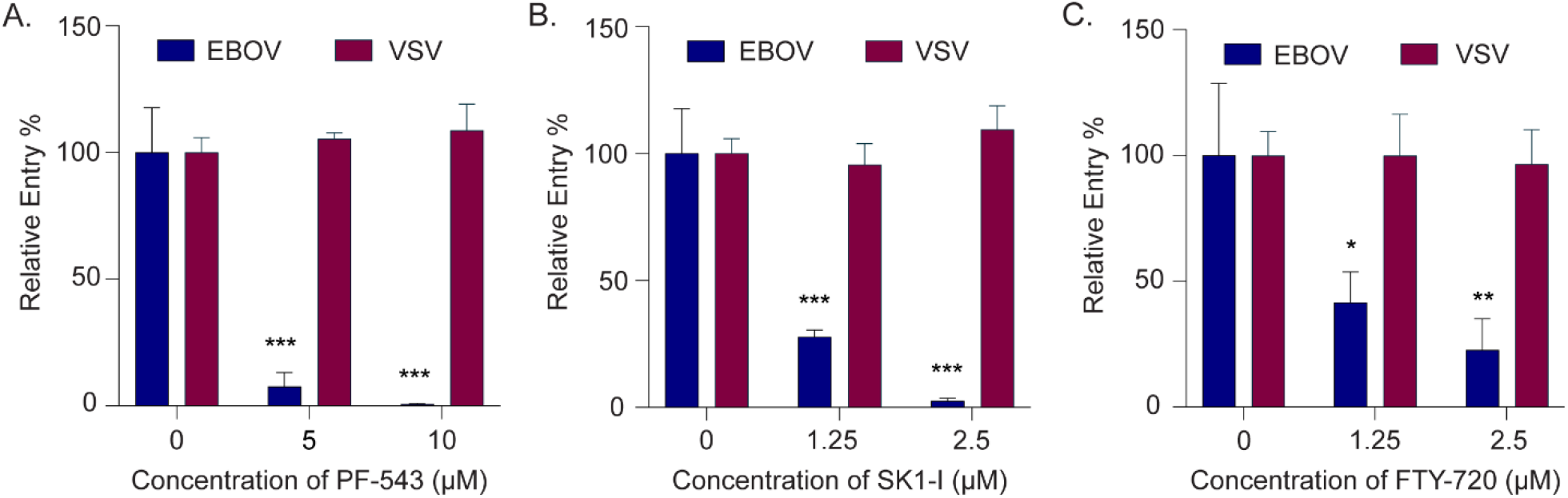
Sphingosine kinase inhibitors block EBOV GP-mediate entry. Entry of βlam VLPs harbouring EBOV GP or VSV G in HT1080 cells treated with (A) PF-543, (B) SK1-I, (C) FTY-720, or vehicle (DMSO, 0.1%) at the indicated concentrations. Relative entry % was determined by measuring the percentage of inhibitor-treated cells with cleaved βlam substrate (CCF2) compared to vehicle-treated cells. Results are expressed as mean ± s.d. of triplicates and are representative of three experiments. Variance for each virus-inhibitor treatment was analyzed by one-way ANOVA, followed by Dunnett’s multiple comparisons test to determine significance. * p < 0.05, ** p < 0.01, *** p < 0.001.

Given their common therapeutic targets, the antiviral activity of PF-543, SK1-I and FTY720 support a role for SKs in EBOV entry. To further confirm this, we synthesized PF-543 analogs with various level of inhibitory activity against SK1, SK2, or both, using a previously published structure-activity-relationship study of the compound (see supporting information for details) (46). The analogs were analyzed for *in vitro* inhibition of SK1 and SK2 activity and for antiviral activity (Table 1, Figure 2). We found that the most potent antiviral compounds were those that strongly inhibited SK1 activity, suggesting that inhibition of EBOV GP-mediated entry is associated with SK1 inhibitory activity (Figure 2). In addition, antiviral activity was severely compromised for compounds that poorly inhibited SK1 activity (>40% residual activity at 1 μM of inhibitor) (Table 1, Figure 2). Lastly, the only compound that was more specific for SK2 and less active against SK1 (Compound 39, SK1 activity: ∼52.6%, SK2 activity: 16.3%) had no antiviral activity (Table 1). Although more work needs to be done to validate the *in cellulo* SK inhibitory activity of these compounds, this panel of inhibitors and the antiviral activity of SK1-I and FTY-720 strongly support a role for SKs, especially SK1, in EBOV GP-mediated entry.

**Table 1.**
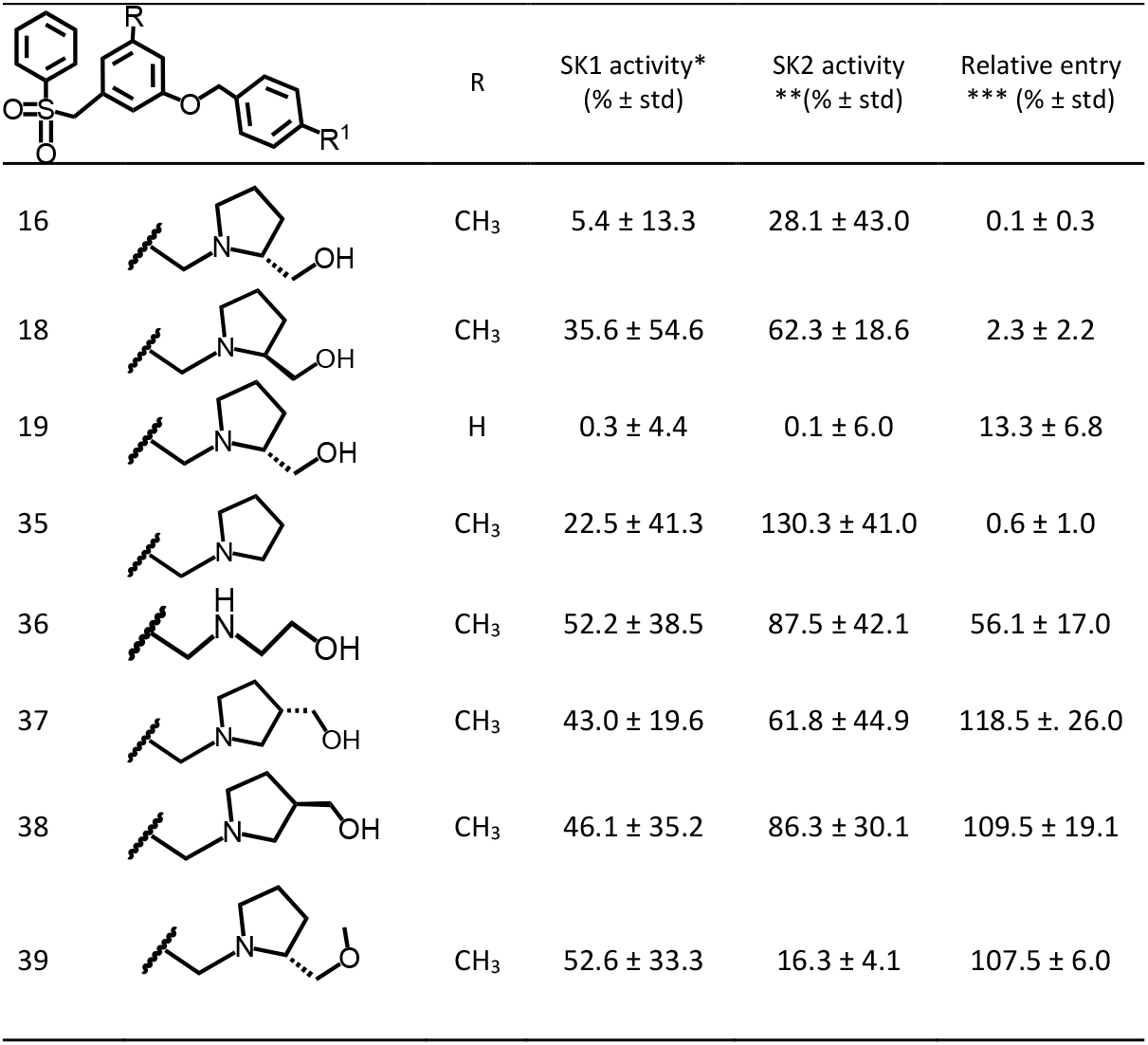
Activity of PF-543 derivative on SK1/2 and EBOV entry.

| | | R | SK1 activity*<br>(% ± std) | SK2 activity<br>**(% ± std) | Relative entry<br>*** (% ± std) |
| --- | --- | --- | --- | --- | --- |
| 16 | | CH<sub>3</sub> | 5.4 ± 13.3 | 28.1 ± 43.0 | 0.1 ± 0.3 |
| 18 | | CH<sub>3</sub> | 35.6 ± 54.6 | 62.3 ± 18.6 | 2.3 ± 2.2 |
| 19 | | H | 0.3 ± 4.4 | 0.1 ± 6.0 | 13.3 ± 6.8 |
| 35 | | CH<sub>3</sub> | 22.5 ± 41.3 | 130.3 ± 41.0 | 0.6 ± 1.0 |
| 36 | | CH<sub>3</sub> | 52.2 ± 38.5 | 87.5 ± 42.1 | 56.1 ± 17.0 |
| 37 | | CH<sub>3</sub> | 43.0 ± 19.6 | 61.8 ± 44.9 | 118.5 ± 26.0 |
| 38 | | CH<sub>3</sub> | 46.1 ± 35.2 | 86.3 ± 30.1 | 109.5 ± 19.1 |
| 39 | | CH<sub>3</sub> | 52.6 ± 33.3 | 16.3 ± 4.1 | 107.5 ± 6.0 |

**Figure 2.**
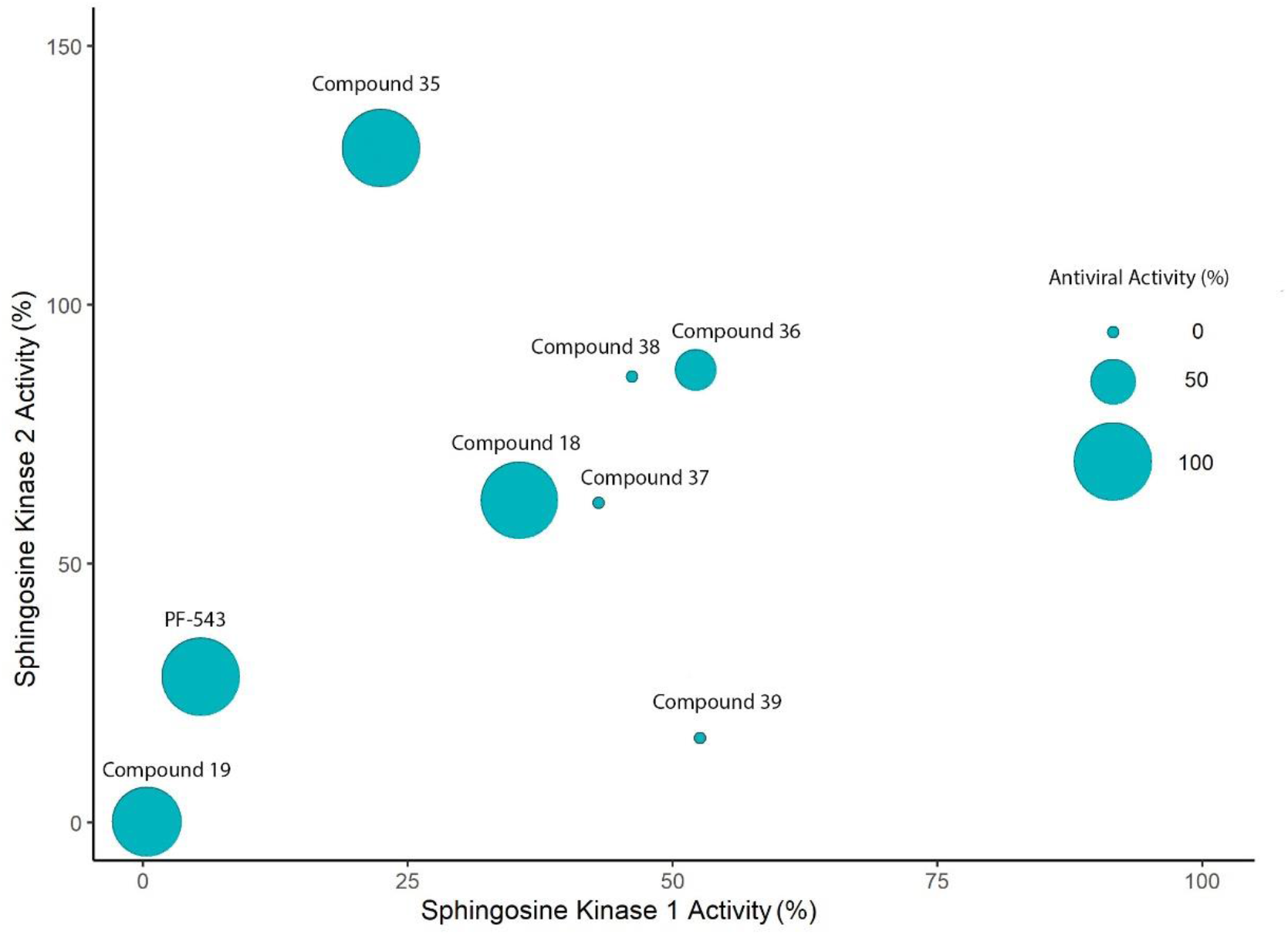
Study of the relationship between antiviral and SK1/2 inhibitory activities using derivatives of PF-543. SK1 and SK2 activity was measured *in vitro* by measuring fluorescence after incubation of PF-543 derivatives (1 µM) or vehicle (DMSO, 0.1%) with SK reaction buffers containing light-sensitive NBD-sphingosine. Percent SK activity was calculated in comparison to vehicle, which represents 100% activity. In parallel, antiviral activity was determined by treating HT1080 cells with the PF-543 derivatives (10 µM) or vehicle (0.1%) and transducing them with MLV pseudotypes encoding LacZ and harbouring EBOV GP. Relative antiviral activity (%) was determined by quantifying LacZ positive inhibitor treated cells compared to vehicle treated cells. Mean SK1 activity (x-axis) and SK2 activity (y-axis) was plotted as a bubble chart using R studio with the size of the bubble representing mean antiviral activity.

### Knockdown of sphingosine kinase 1 and 2 reduces EBOV GP-mediated entry

To directly assess a role for SKs, we targeted SK1 and SK2 using Dicer-Substrate Short Interfering RNAs (DsiRNAs). SK1 or SK2 targeting DsiRNAs, or a non-targeting negative control, were transfected in HT1080 cells. Immunoblot analysis confirmed that SK1 and SK2 expression was substantially knocked down when the respective DsiRNA was transfected (Figure 3A). VLPs harboring EBOV GP or VSV-G were then used to measure entry (Figure 3B). While knockdown of SK1 or SK2 did not interfere with VSV-G-mediated entry, a decrease in entry of EBOV VLPs was observed. Interestingly, although expression of each SK was undetectable when cells were transfected with their respective targeting DsiRNA, SK2 knockdown had a slightly greater inhibitory effect on EBOV entry (Figure 3B). To test whether they play redundant roles in viral entry, we tried to perform double knockdown. Unfortunately, viability of the cells was severely compromised (data not shown). Nevertheless, these results suggest that both SK1/2 play a role in infection, providing even further evidence that EBOV GP-mediated entry requires SK1 and/or SK2 activity.

**Figure 3.**
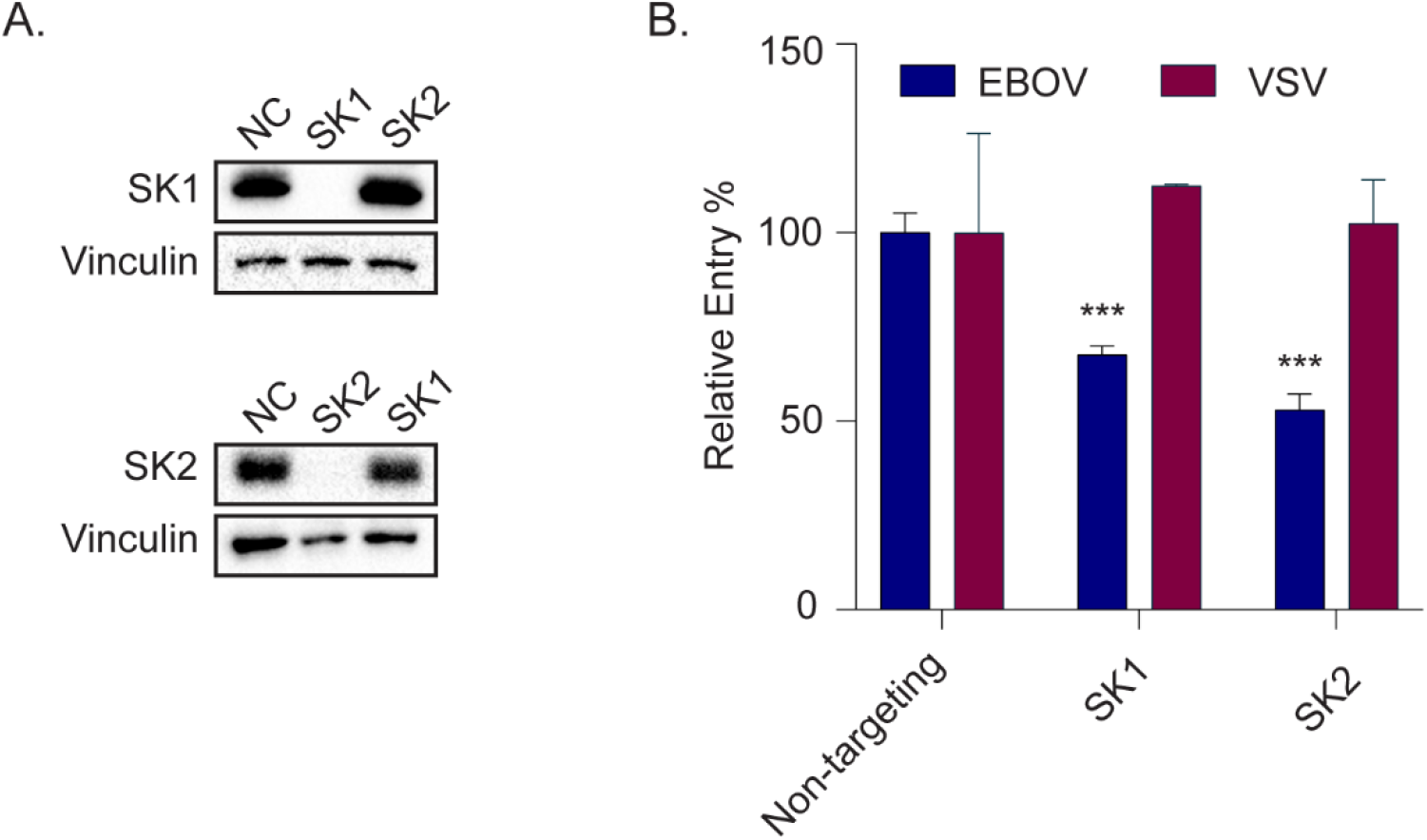
SK1/2 knockdown blocks entry of EBOV in HT1080. HT1080 cells transfected with nontargeting control dsiRNA (NC: negative control) or dsiRNA targeting SK1 or SK2 were either (A) lysed and expression of SK1 or SK2 (upper) and Vinculin (lower) detected by immunoblot, or (B) infected with βlam VLPs harbouring EBOV GP or VSV G. Relative entry % was determined by measuring the percentage of inhibitor-treated cells with cleaved βlam substrate (CCF2) compared to vehicle-treated cells. Results are expressed as mean ± s.d. of triplicates and are representative of three experiments. Variance was analyzed by one-way ANOVA, followed by Dunnett’s multiple comparisons test to determine significance. *** p < 0.001.

### PF-543 blocks EBOV growth

We next sought to confirm that SK1/2 activity is required for replicative EBOV infection. Given that our data indicate that both SK1 and SK2 are involved in entry, we chose to investigate the effect of PF-543, which inhibits both SKs, on infection by replication-competent EBOV. Vero cells were infected with Green Fluorescent Protein (GFP)-expressing replicative EBOV in the presence or absence of PF-543 and growth was assessed by measuring GFP fluorescence. We found that PF-543 significantly inhibited replicative EBOV at 5µM and completely prevented it at 10µM (Figure 4 AB). These results further support a role of SKs in EBOV infection and suggest that they are potential targets for antiviral therapy.

**Figure 4.**
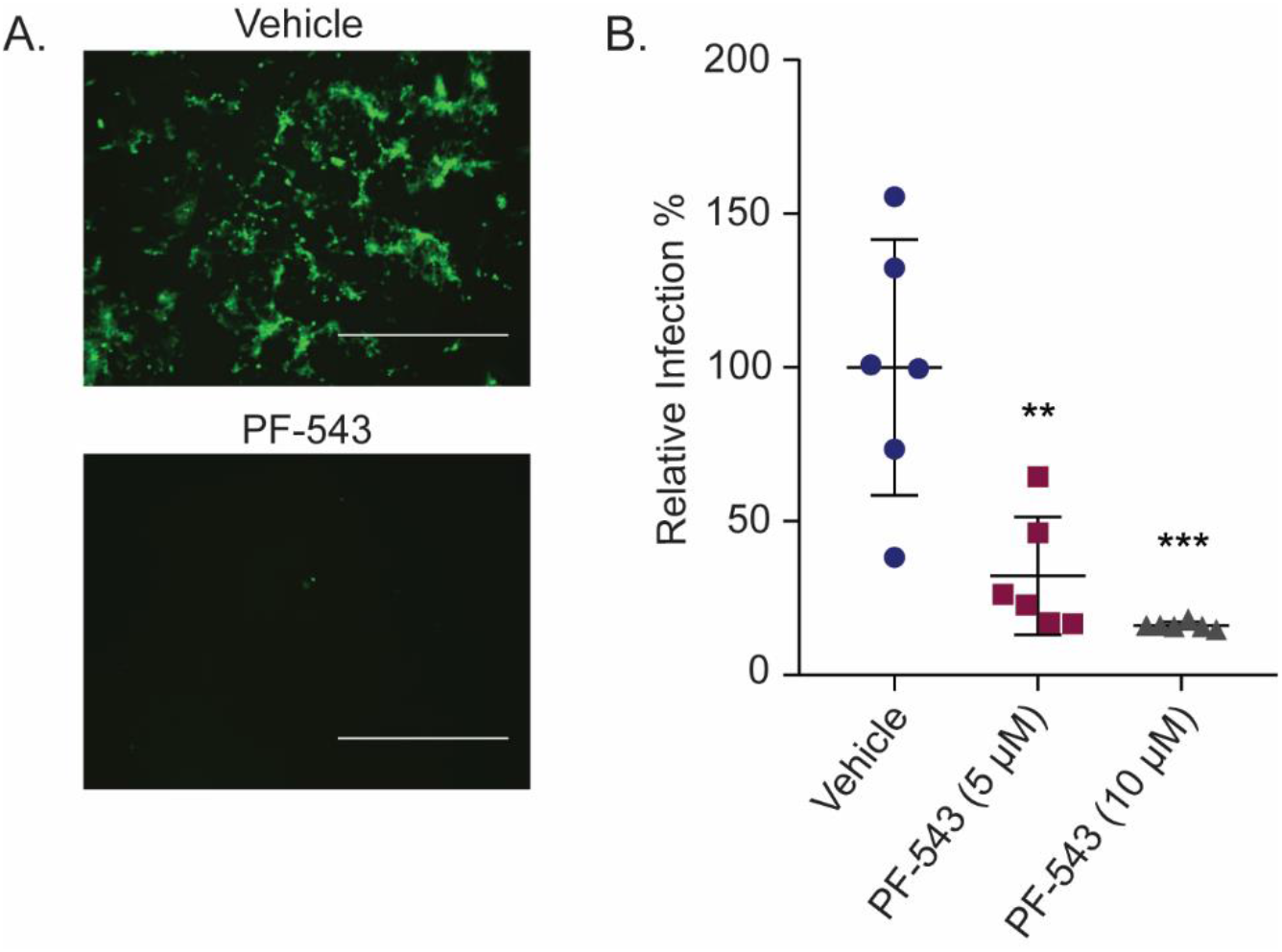
PF-543 blocks infection of replication-competent EBOV. Infection of Vero cells with replication-competent EBOV expressing GFP and treated with vehicle or PF-543. Images in (A) were acquired for vehicle or PF-543 (5 µM) treated wells, bar=1mm. Relative infection % in (B) was determined by measuring mean GFP fluorescence intensity for PF-543 treated cells compared to vehicle alone. Results are expressed as mean ± s.d. of three experiments. Variance was analyzed by one-way ANOVA, followed by Dunnett’s multiple comparisons test to determine significance. * p < 0.05, ** p < 0.01, *** p < 0.001.

### Sphingosine kinase inhibitors do not interfere with EBOV attachment and internalization

EBOV entry begins with attachment of the viral particle to the cell surface, which is mediated by multiple cellular attachment factors, followed by virion internalization via a macropinocytosis-like mechanism (10). To further elucidate the role of sphingosine kinases in EBOV entry, we first assessed whether the SK inhibitors reduced attachment of EBOV using EBOV VLPs containing VP40 fused to GFP. We exposed inhibitor-treated cells to EBOV VLPs at 4 °C to only allow for virion attachment and assessed the resulting VLP fluorescence by flow cytometry. We found that none of the inhibitor treatments resulted in a reduction in VLP fluorescence, indicating that SK inhibitors do not significantly block attachment of EBOV VLPs (Figure 5A).

**Figure 5.**
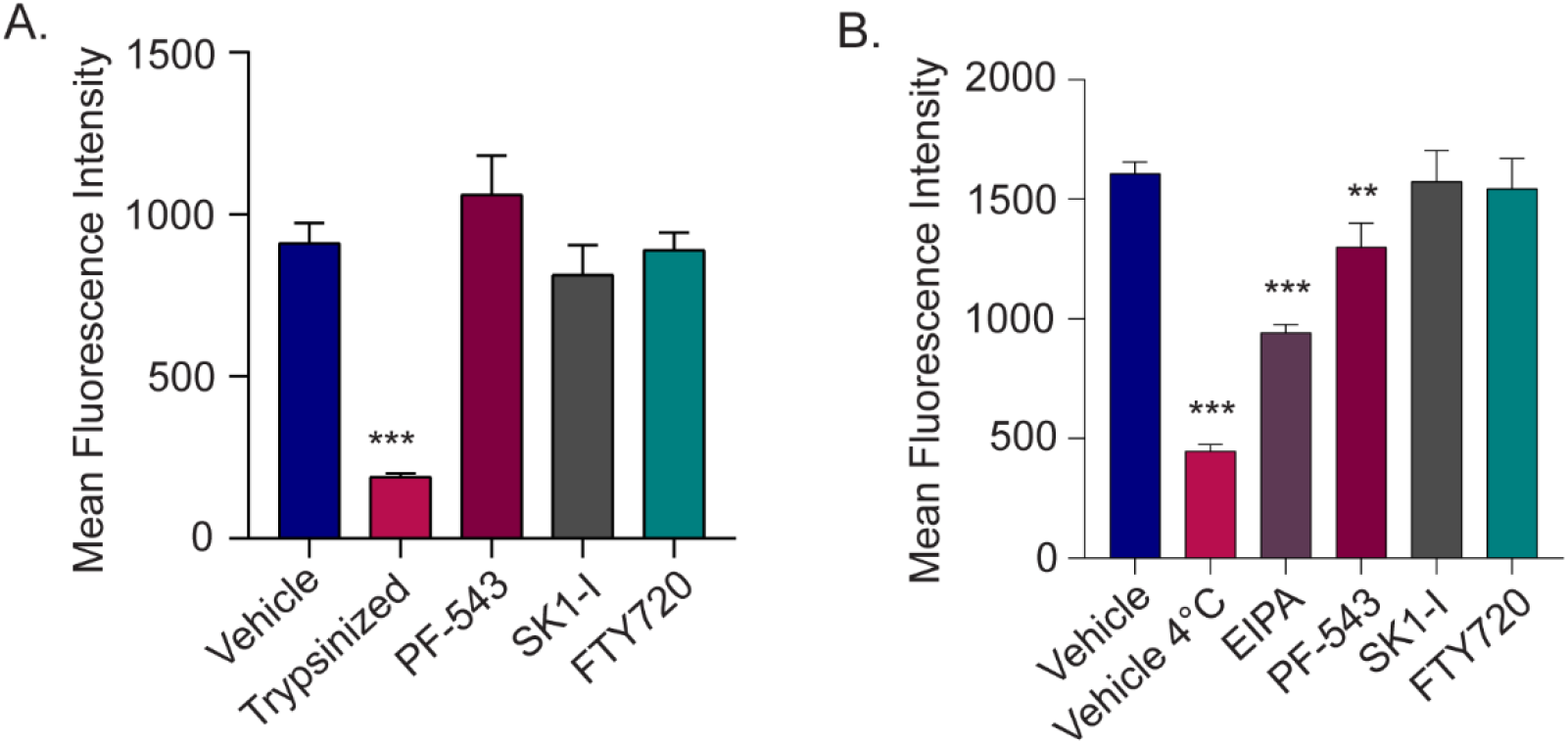
SK inhibitors do not interfere with attachment and have minimal or no effect on internalization. HT1080 cells were pre-treated with vehicle (DMSO, 0.1%), PF-543 (10 µM), SK1-I (2.5 µM), FTY720 (2.5 µM), or EIPA (30 µM) for 1 hour. In (A), cells were then detached and incubated with GFP-VLPs harbouring EBOV GFP at 4 ºC. Cells were washed to remove unbound VLPs and a subset of cells were trypsinized to remove bound VLPs as a negative control. Cells were incubated with SYTOX red dead cell stain prior to analysis of fluorescence by flow cytometry. (B) GFP-VLPs harbouring EBOV GP were pre-bound to treated cells by spinoculation at 4 ºC, washed, and either incubated again with vehicle or inhibitor at 37 ºC, or vehicle alone at 4 ºC. Cells were then trypsinized to remove bound VLPs, washed, and treated with SYTOX red dead cell stain prior to analysis of fluorescence by flow cytometry. Results are expressed as mean ± s.d. and are representative of three experiments. Variance was analyzed by one-way ANOVA, followed by Dunnett’s multiple comparisons test to determine significance. ** p < 0.01, *** p < 0.001.

To assess if the SK inhibitors blocked macropinocytic uptake of EBOV, we allowed virus internalization at 37 °C following the 4 °C attachment step. As controls, some samples were kept at 4°C throughout the experiment to ensure that internalization could not occur, and some samples were incubated with 5-(*N*-Ethyl-*N*-isopropyl)amiloride (EIPA), a known macropinocytosis inhibitor (Figure 5B) (47). We found that while PF-543 had a modest effect on internalization of fluorescent EBOV VLPs, the effect was not nearly as pronounced as that of EIPA (Figure 5B). Notably, SK1-I and FTY720 had no effect on EBOV VLP internalization (Figure 5B). Taken together, these results suggested that SK activity is required at an entry step after attachment and internalization.

### Entry of pre-cleaved EBOV VLPs is inhibited by sphingosine kinase inhibitors

A critical post-internalization EBOV entry step is the cleavage of EBOV GP by endosomal cathepsin proteases such as cathepsin B (1). Interestingly, a previous study suggested that SK1-I treatment results in enlarged vacuoles that lack cathepsin B activity, therefore we also assessed if the SK inhibitors may prevent GP-mediated cleavage by cathepsins (27). To investigate this, we experimentally mimicked cathepsin cleavage by pre-cleaving EBOV VLPs with thermolysin. We assessed entry of uncleaved and cleaved VLPs in HT1080 cells treated with a cathepsin B inhibitor, Ca074, in addition to the SK inhibitors. As expected, Ca074 had reduced potency on cleaved VLPs compared to the uncleaved VLPs (Figure 6). In contrast, the SK inhibitors were equally potent at inhibiting entry of the cleaved and uncleaved VLPs, suggesting that their primary mechanism of action is not to inhibit Cathepsin B-mediated cleavage of EBOV GP.

**Figure 6.**
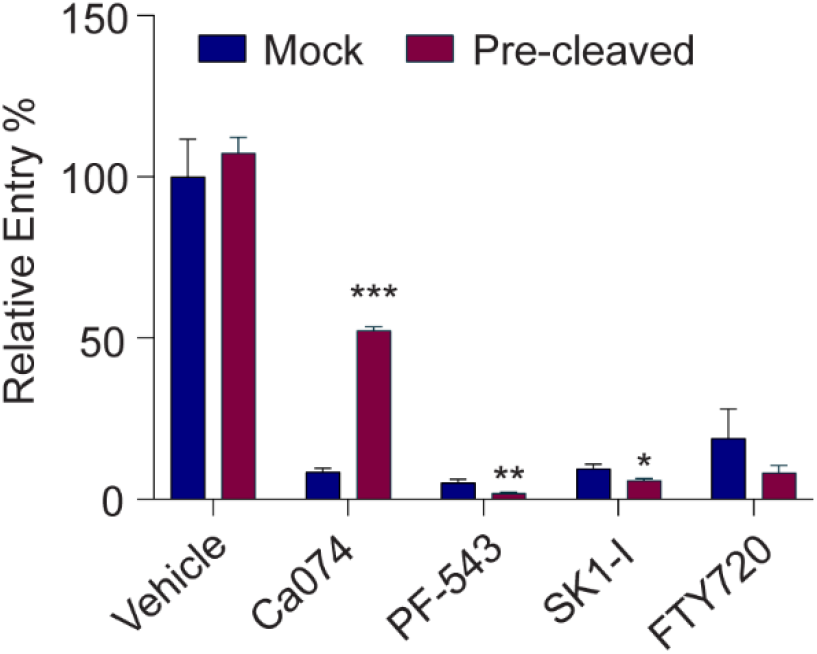
Pre-cleaving EBOV GP does not rescue the SK inhibitor entry block. HT1080 cells were pre-treated with vehicle (DMSO, 0.1%), Ca074-Me (2.5 µM), PF-543 (10 µM), SK1-I (2.5 µM), or FTY720 (2.5 µM) for 1h prior to incubation with βlam VLPs harbouring either thermolysin cleaved (pre-cleaved) or mock cleaved EBOV GP. Relative entry % was determined by measuring the percentage of inhibitor-treated cells with cleaved βlam substrate (CCF2) compared to vehicle-treated cells. Results are expressed as mean ± s.d. and are representative of three experiments. Unpaired t-tests were performed to determine significance between mock and pre-cleaved samples per inhibitor treatment. * p < 0.05, ** p < 0.01, *** p < 0.001.

### Sphingosine kinase inhibitors interfere with trafficking of EBOV to NPC1+ intracellular compartments

Since the SK inhibitors did not significantly prevent attachment and internalization, we next sought to assess whether they blocked virus endolysosomal trafficking to the EBOV receptor, NPC1. For these experiments, we utilized fluorescent VLPs harbouring EBOV GP^F535R^, which can bind to NPC1 following cathepsin cleavage but cannot undergo fusion, therefore allowing us to visualize accumulation of VLPs in NPC1+ compartments (5). As a control, we used Akt Inhibitor VIII, which has been previously shown to result in accumulation of EBOV in early endosomal compartments (17, 18). We determined the percentage of fluorescent VLPs colocalizing with immunostained NPC1 in inhibitor treated cells compared to vehicle alone and found a decrease in the percentage of VLPs per cell that colocalized with NPC1 for all three SK inhibitors (Figure 7AB), suggesting that the SK inhibitors interfere with endosomal trafficking. We also observed the presence of dilated intracellular vesicles in SK1-I and FTY720 treated cells that were not present in vehicle or PF-543 treated cells (Figure 7A, red arrows). This is consistent with recent studies demonstrating that treatment with sphingosine or sphingosine analogs, including SK1-I and FTY720, induce rapid formation of enlarged intracellular vesicles in cells (26, 27). Since such vesicles are often indicative of defects in endosomal trafficking, their presence also supports a model by which treatment with the SK inhibitors result in endosomal trafficking defects.

**Figure 7.**
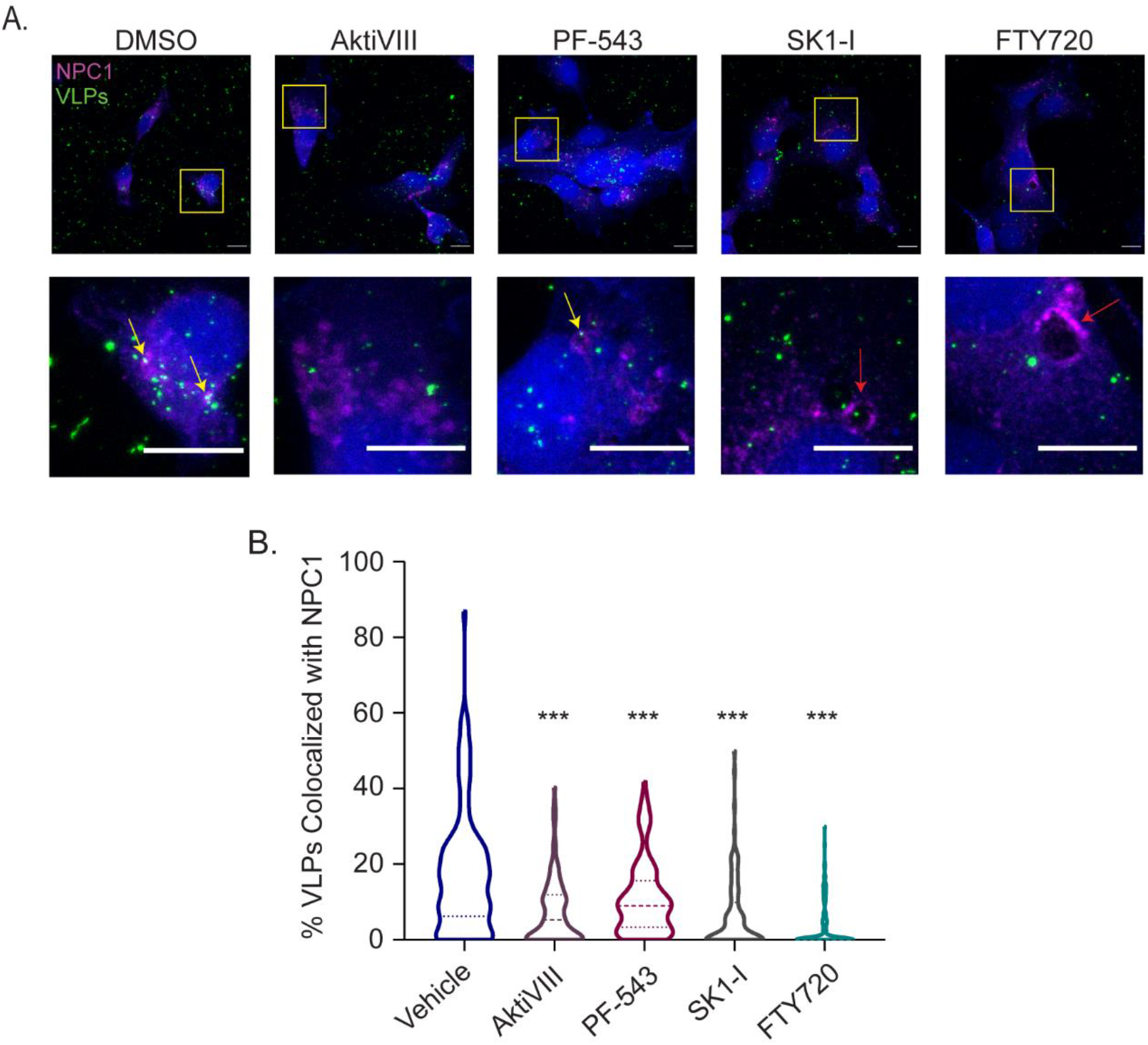
SK inhibitors block trafficking of EBOV VLPs to NPC1. (A) HT1080 cells were pre-treated with vehicle (DMSO, 0.1%), AktiVIII (10 µM), PF-543 (10 µM), SK1-I (2.5 µM), or FTY720 (2.5 µM) for 1h, followed by incubation with GFP-VLPs (Green) harbouring the fusion deficient ΔM GPF535R for 3h. After 2.5h, CMAC cytoplasmic dye (Blue) was added to the media for the remaining 30 minutes of the incubation. Cells were then fixed, permeabilized, and immunostained with anti-NPC1 and DY-650 conjugated antiserum (Magenta). Cells were imaged on a LSM800 confocal microscope (Zeiss). Displayed images are maximum intensity Z-projections, yellow arrows indicate colocalization of VLPs and NPC1, red arrows point to dilated intracellular vesicles, bar = 10 µm. (B) Colocalization between VLPs and NPC1 was analyzed using Imaris software (Bitplane). Results are expressed as mean ± s.d. and are of three experiments. Variance was analyzed by one-way ANOVA, followed by Dunnett’s multiple comparisons test to determine significance. *** p < 0.001.

### The endolysosomal trafficking defect induced by the SK inhibitors is independent of S1P Receptor (S1PR) signaling

S1P, the product of SKs, is a bioactive sphingolipid involved in regulating cell growth, survival, and migration (48). It is well-characterized for binding to cell surface S1PRs and activating signaling cascades such as the PI3K-Akt pathway (23, 49, 50). Since Akt signaling was shown to be important for EBOV trafficking and given that SK inhibitors were previously shown to inhibit Akt phosphorylation, we hypothesized that the SK inhibitors block EBOV trafficking by preventing S1P activation of S1PRs, therefore preventing subsequent Akt activation (16-18, 51). To test this hypothesis and dissect the effect of SKs and S1PR signalling, we evaluated the antiviral activity of FTY-720-Phosphate, which inhibits S1PR signaling by inducing S1PR endocytosis and degradation but, unlike FTY-720, cannot enter the cell and therefore does not inhibit SKs (43, 52, 53). Cells were pre-treated for one hour with FTY-720 or FTY-720-Phosphate and entry of VLPs harbouring EBOV GP or VSV G was evaluated. We found that FTY-720-Phosphate had no effect on EBOV-GP mediated entry while FTY-720 drastically blocked entry at the same concentration (Figure 8A), suggesting that the trafficking defect caused by SK inhibitors is not a result of reduced signaling through S1PRs.

**Figure 8.**
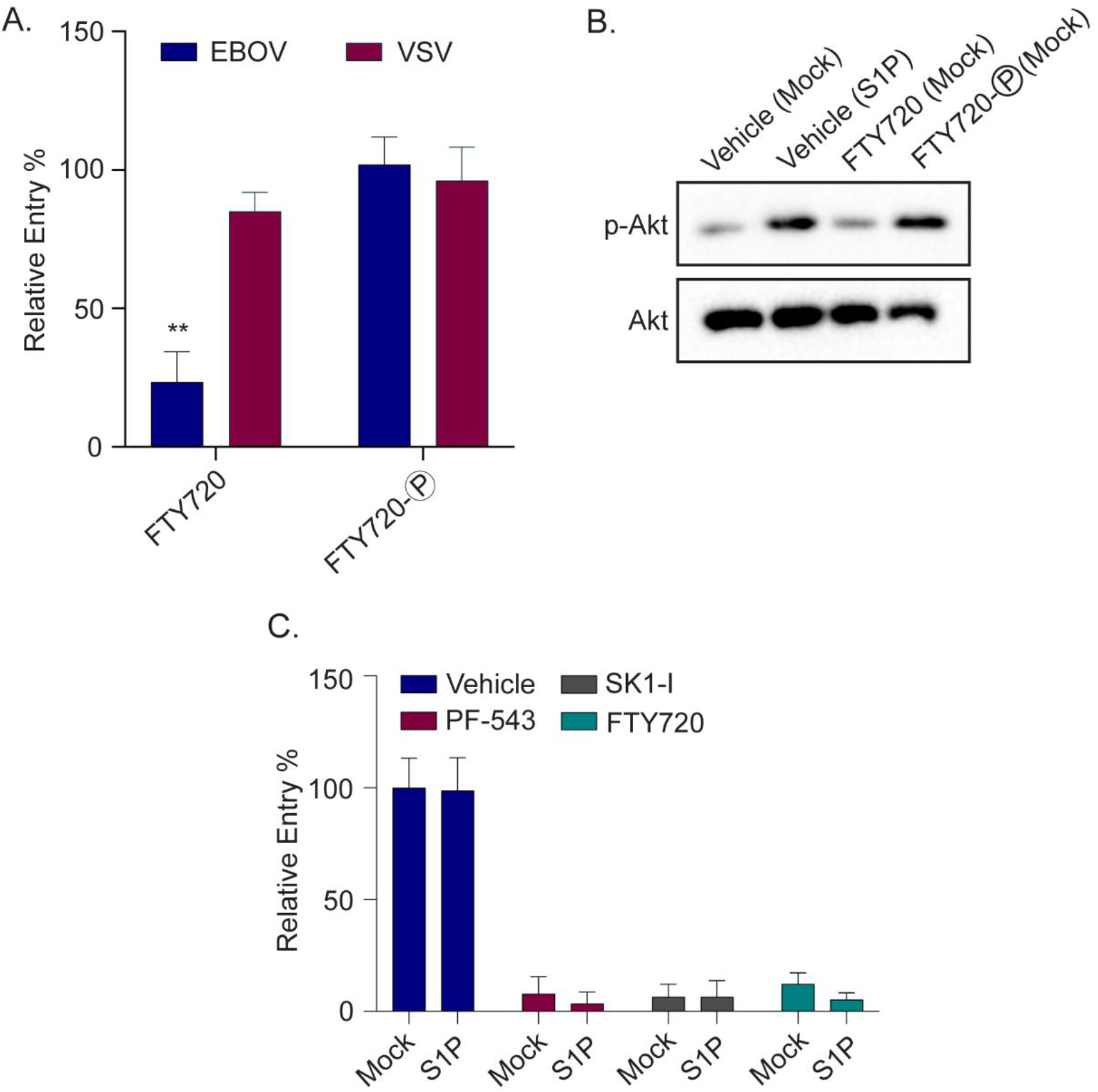
S1PR agonist does not reduce EBOV entry and S1PR activation by S1P does not rescue EBOV infection in inhibitor treated cells. (A-B) HT1080 cells were pre-treated with vehicle (DMSO, 0.1%), FTY720 (2.5 µM), or FTY720-Phosphate (2.5 µM) for 1h prior to (A) infection with βlam VLPs harbouring EBOV GP or VSV G, or (B) stimulation with S1P (125 nM) or mock. In (A), relative entry % was determined by measuring the percentage of inhibitor-treated cells with cleaved βlam substrate (CCF2) compared to vehicle-treated cells. In (B), cells were lysed and immunoblotted for phosphorylated Akt (p-Akt S473) or total Akt. (C) HT1080 cells were pre-treated with vehicle (DMSO, 0.1%), PF-543 (10 µM), SK1-I (2.5 µM), or FTY720 (2.5 µM), stimulated with S1P (125 nM) or mock, and infected with βlam VLPs harbouring EBOV GP. Relative entry % was determined by measuring the percentage of inhibitor-treated cells with cleaved βlam substrate (CCF2) compared to vehicle-treated cells. Results are expressed as mean ± s.d. of triplicates and are representative of three experiments. Unpaired t-tests were performed to determine significance between treatments. ** p < 0.01.

To further explore if FTY-720 and FTY-720-Phosphate reduce S1PR mediated PI3K-Akt pathway activation, we treated cells with the inhibitors and assessed Akt phosphorylation by immunoblot. As a positive control, we added S1P to vehicle-treated cells and found that S1P treatment increased the levels of phosphorylated Akt in cells. Interestingly, while treatment with FTY-720 did not alter phosphorylated Akt levels compared to vehicle treated cells, treatment with FTY-720-Phosphate increased Akt phosphorylation to a similar extent as S1P treated cells (Figure 8B). This suggests that FTY-720-Phosphate induces activation of S1PRs prior to inducing their degradation, a result that is consistent with previous reports in the literature (53). Therefore, the lack of entry inhibition seen after FTY-720-Phosphate treatment does not fully rule out the possibility that the antiviral activity of the SK inhibitors is due to a reduction in S1P signaling through cell surface S1PRs.

To address this, we performed an add-back experiment where S1P was added to inhibitor treated cells prior to virus addition. If the entry and trafficking defects observed after SK inhibitor treatment are due to a decrease in S1P signaling through S1PRs, adding extracellular S1P should rescue the inhibitory entry effects mediated by the SK inhibitors. Importantly, we found that S1P addition did not rescue EBOV entry in the presence of PF-543, SK1-I, or FTY720 (Figure 8C). This result strongly suggests that the trafficking defect observed after treatment with the SK inhibitors is not due to reduced S1P signaling through S1PRs. Further work needs to be done to elucidate the precise molecular mechanism by which inhibiting SKs results in an endosomal trafficking defect that blocks EBOV entry.

### Sphingosine kinase inhibitors reduce entry of other late penetrating viruses

The host factors important for EBOV intracellular trafficking may be shared with other viruses whose entry receptor or other triggering factors are also localized in late endosomes. As such, it is plausible that entry mediated by the GP of these viruses may also be inhibited by the SK inhibitors. Therefore, we tested the ability of the SK inhibitors to block entry of a panel of enveloped viruses that require various extents of endosomal trafficking in all or some of the cell types that they infect. Specifically, we tested Marburg virus (MARV), a related filovirus that shares its entry receptor with EBOV, as well as Junin virus (JUNV), Influenza A virus (IAV), and LFV, all of which require low pH for entry, while LFV further requires the lysosomal protein LAMP-1 (2, 3, 28, 29, 31, 54-58). As controls, we included VSV, which undergoes fusion in early endosomes, and Nipah virus (NiV), which requires its entry receptor, ephrinB2, to mediate fusion at the cell surface (59-62). To explore the requirement of SKs for entry of these viruses, we utilized MLV pseudotypes harbouring their respective viral fusion proteins and tested entry in the presence of the SK inhibitors. While the SK inhibitors were the most potent at inhibiting EBOV entry, they also significantly inhibited MARV entry, and to a lesser extent LFV and JUNV (Figure 9A). Unexpectedly, these inhibitors either enhanced IAV entry, as was the case for PF-543 and SK1-I, or had no effect, as was the case for FTY720 (Figure 9A). Lastly, we found that PF-543 and SK1-I did not reduce VSV or NiV entry, and while FTY720 slightly reduced VSV entry, it was not to the same extent as the other viruses (Figure 9A).

**Figure 9.**
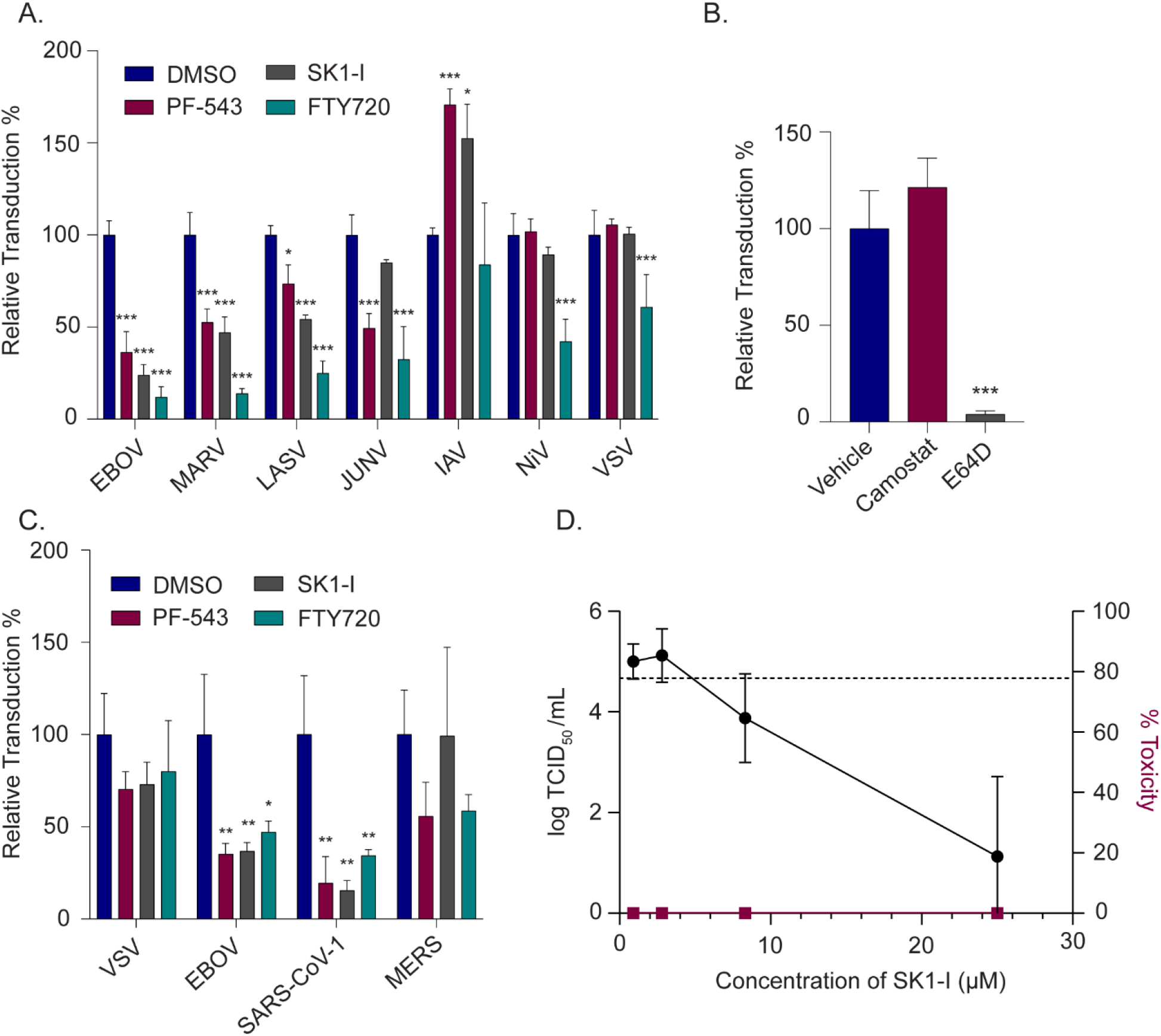
Sphingosine kinase inhibitors inhibit entry of late penetrating viruses and SARS-CoV-2 infection of Huh7.5 cells. (A-C) HT1080 cells treated with vehicle (DMSO, 0.1%), PF-543 (10 µM), SK1-I (2.5 µM), FTY720 (2.5 µM), Camostat (25 µM), or E64D (10 µM) for 1h were transduced with (A) MLV pseudotypes or (B,C) lentiviral pseudotypes, encoding lacZ and harbouring the fusion proteins of the indicated viruses. Relative transduction % was determined by quantifying LacZ positive inhibitor treated cells compared to vehicle treated cells. (D) Huh7.5 cells treated with vehicle or increasing concentrations of SK1-I were infected with replication-competent SARS-CoV-2. 48 hpi, supernatant was collected and used directly to measure the TCID50. Results are expressed as mean ± s.d. of triplicates and are representative of three experiments. Variance was analyzed by one-way ANOVA, followed by Dunnett’s multiple comparisons test to determine significance. * p < 0.05, ** p < 0.01, *** p < 0.001.

In addition to the aforementioned viruses, many coronaviruses also require triggering factors in late endosomes to mediate entry in some cell types. Therefore, we utilized lentivirus pseudotypes to examine entry mediated by the spike protein of the coronaviruses SARS-CoV-1 and middle eastern respiratory syndrome coronaviruses (MERS) in HT1080 cells. Importantly, in these cells, SARS-CoV-1 S-mediated entry is dependent on cathepsin activity (Figure 9B), indicating a requirement for late endosomal trafficking. Accordingly, we found that all SK inhibitors drastically inhibited SARS-CoV-1 entry, while only PF-543 had a significant effect for MERS entry (Figure 9C). Lastly, we tested whether the most potent inhibitor for inhibiting SARS-CoV-1 entry, SK1-I, also inhibited infection of replication competent SARS-CoV-2. We found that SK1-I inhibited infection of SARS-CoV-2 in Huh7.5 cells in a dose-dependent manner (Fig. 9D). Taken together, these results suggest that sphingosine kinases are important host factors that help facilitate viral entry of a range of enveloped viruses that enter cells through endosomes and therefore could serve as host targets for novel pan-viral antiviral therapies.

## Discussion

Entry of enveloped viruses into host cells requires fusion between the viral and host cell membranes, a process mediated by viral fusion proteins. For EBOV and other viruses, cellular proteins important for the priming and/or triggering of viral fusion proteins are localized in late-endosomes and lysosomes. Therefore, cellular entry of these viruses requires internalization and endolysosomal trafficking. In this study, we investigated SK inhibitors and characterized the role of SKs in EBOV entry. We found that SKs were required for EBOV trafficking to late intracellular vesicles containing the viral receptor and, accordingly, SK inhibitors also blocked infection by other viruses that use late endosomes/lysosomes for membrane fusion and entry. Our studies highlight an important role for SKs in endosomal trafficking and suggest that SK inhibitors have the potential to be developed for pan-antiviral therapy.

Signaling lipids play critical roles in endocytosis and endocytic trafficking (63, 64) and lipid modifying enzymes have been implicated to be important for EBOV entry; these include PIKfyve, diacylglycerol kinases (DGKs), and acid sphingomyelinases (ASMase) (13, 16, 65). For instance, EBOV was found to associate with lipid rafts rich in sphingomyelin (SM) and ASMases, and knockdown of ASMase inhibited EBOV entry (65). ASMase hydrolyses SM to ceramide, which can be converted to sphingosine and subsequently S1P via SKs. Ceramide has been implicated in lipid raft expansion and is more favorable compared to SM to promote membrane curvature and macropinocytosis (66). Interestingly, previous studies have demonstrated that SK1 associates with membranes of high curvature including macropinosomes and speculate that endocytosis may correlate with the conversion of SM to sphingosine, which can flip across the membrane and be converted to S1P by SKs (67). In our study, we also found that treatment with the SK1 inhibitor, PF-543, slightly reduced internalization of EBOV, suggesting that SK1 may play a small role in macropinocytic uptake of EBOV (Figure 4B). Interestingly, another study showed that a SK1 activator, K6PC-5, inhibited EBOV GP-mediated entry (21). Therefore, taken with our current study, this suggests that interfering with S1P production, whether it be positively or negatively, leads to inefficient EBOV entry. Although the authors did not investigate the specific entry step that is inhibited in the presence of K6PC-5, they found that the antiviral activity was independent of S1P receptors, which is also congruent with our findings (21). The specific role(s) of S1P during entry and whether the lipid composition of cellular membranes at the site of EBOV attachment or internalisation changes remains to be investigated.

Here we show that all SK inhibitors prevented EBOV from trafficking to late-endosomes and/or lysosomes containing NPC1 (Figure 4) suggesting that SKs play critical roles in endosomal trafficking. Interestingly, a few other studies have also implicated SKs to be important for proper endosomal maturation and fusion (26, 27). In a previous study, treatment with the sphingosine analogues SK1-I and FTY720 resulted in the formation of dilated intracellular vesicles that are positive for late endosomal markers Rab7a and LAMP-1, suggesting that the trafficking block observed for EBOV after treatment with these inhibitors occurs at the late endosome step (26, 27). Furthermore, Young et al. found that the vacuolization is independent of S1P signaling, as pre-treatment with FTY720-Phosphate did not alter the SK1-I/FTY720 induced vacuole formation (27). Moreover, they show that overloading endosomal membranes with sphingosine results in similar vacuole formation and that treatment with PF-543 delays clearance (27). Our data indicates that while PF-543, a SK inhibitor that is not a sphingosine analogue, does not induce formation of dilated vesicles, it does induce a similar endocytic trafficking defect (Figure 4C and D). This not only provides an explanation for the mechanism of antiviral activity of our SK inhibitors but also points to some potential areas for future investigations of the role of sphingosine, SKs, and S1P in the regulation of endolysosomal trafficking.

In addition to blocking EBOV entry, the SK inhibitors tested also inhibited entry of other late penetrating viruses. For example, MARV, a related filovirus, was sensitive to the inhibitory effects of the SK inhibitors, although not to the same extent as EBOV (Figure 5A). Similarly, LFV entry was slightly blocked by the SK inhibitors, but was significantly less sensitive in comparison to EBOV (Figure 5A). These differences may be attributed to the ability of both MARV, and even more so, LFV, to undergo fusion in earlier endocytic compartments (4, 58). Studies have shown that although MARV does require NPC1 for fusion, the MARV GP is significantly less stable than EBOV GP and although proteolytic triggering of MARV GP is required for entry, MARV is less dependent on cathepsins B and L (4, 68, 69). This suggests that MARV GP-mediated fusion may be able to occur in earlier endocytic compartments compared to EBOV. Similarly, a recent study suggested that LFV fusion may be able to occur in early endosomes that contain only a small amount of the entry receptor LAMP-1 (58). We also tested the effect of the SK inhibitors on coronavirus entry. For coronaviruses, the site of entry depends on the expression of host proteases required to cleave the spike protein for activation (36). In our study, we tested the effects of the SK inhibitors in cell types lacking surface proteases to allow for entry via a route that requires endosomal trafficking. We found that the SK inhibitors inhibited entry and/or infection by SARS-CoV-1, SARS-CoV-2, and to a lesser extent, MERS (Figure 9). Together, our data strongly suggests that the SK inhibitors block a late trafficking step, and that the efficacy of these inhibitors as antiviral agents will depend on the location of fusion triggering among different viruses.

In this study, we identified sphingosine kinases as host factors required for viral entry of EBOV and that of other late-penetrating viruses. We found that inhibition of SKs impairs endosomal trafficking and that this occurs independently of S1P signaling through S1PRs at the cell surface. Importantly, since entry of many enveloped viruses requires endosomal trafficking, small molecule inhibitors of SKs may be promising candidates for the development of broad-spectrum antiviral therapeutics that could prevent infection of both current and newly emerging viruses.

## Materials and Methods

### Cell lines, antibodies, and inhibitors

HEK293T cells (ATCC) were cultured in Dulbecco’s Modified Eagle Medium (DMEM, Wisent), while HT1080 and Vero cells (ATCC) were cultured in Minimum Essential Medium (MEM, Sigma). Both culture media were supplemented with 10% Fetal Bovine Serum (FBS, Sigma), 0.3 mg/mL L-glutamine, 100 U/mL penicillin, and 100 µg/mL streptomycin (Wisent). Cells were maintained at 37 ºC in 5% CO_2_ at 100% relative humidity.

Primary antibodies used were NPC1 (ab134113, Abcam), Akt (9272S, Cell Signaling Technology), phospho-Akt S473 (92721S, Cell Signaling Technology), GAPDH (ab8245, Abcam), SK1 (12071S, Cell Signaling Technology), SK2 (32346S, Cell Signaling Technology) and pan-filovirus anti-GP antibody (21D10, IBT Bioservices). Secondary antibodies used were goat anti-rabbit IgG HRP-linked (7074S, Cell Signaling Technology), anti-mouse IgG HRP-linked (7076S, Cell Signaling Technology) and DY650 sheep anti-rabbit (ab96926, Abcam).

PF-543 (Cayman Chemical), SK1-I (BML-258, Enzo Life Sciences Inc.), FTY720 (Cayman Chemical), FTY720 Phosphate (Cayman Chemical), 5-(N-ethyl-N-isopropyl)-Amiloride (EIPA, Cayman Chemical), and Akt Inhibitor VIII (Cayman Chemical) were prepared in DMSO, aliquoted, and stored at -20 ºC prior to use. PF-543 derivatives were synthesized as described in supplementary information. Sphingosine-1-phosphate (d18:1, Cayman Chemical) was prepared in a methanol:water (95:5), mixture was heated and sonicated for dissolution, and stored in glass vials at -20 ºC. Prior to use, the methanol:water solution was evaporated under a N_2_ stream and a 125 µM stock solution was prepared in 4 mg/mL fatty acid free Bovine Serum Albumin (Sigma) in PBS and stored at -20 ºC for up to 3 months.

### Plasmids and DsiRNA

Plasmids encoding the virus glycoproteins EBOV Δmucin GP, EBOV Δmucin GP^F535R^, MARV GP, VSV-G, LFV GP, NiV F and G fusion proteins, IAV Vict HA and 1918 NA, the MLV packaging plasmid, and MLV retroviral vector encoding LacZ were kind gifts of Dr. James Cunningham (Brigham and Women’s Hospital, Boston, Massachusetts). Plasmids encoding the EBOV NP and EBOV VP40-β-lactamase or GFP were kind gifts of Dr. Lijun Rong, University of Illinois. The plasmids encoding human ACE2 and SARS-CoV-S were a kind gifts of Dr. Hyeryun Choe (The Scripps Research Institute, Jupiter, Florida). Plasmid encoding the MERS-CoV-S was a kind gift of Thomas Gallagher (Loyola University Chicago, Chicago, Illinois). LV-Lac was a gift from Inder Verma (Addgene plasmid # 12108, (70)) and psPAX2 was a gift from Didier Trono (Addgene plasmid # 12260).

Predesigned DsiRNAs for SK1 (hs.Ri.SPHK1.13.1; duplex sequences: 5’-rGrCrGrUrCrArUrGrCrArUrCrUrGrUrUrCrUrArCrGrUrGCG -3’;

3’-rCrGrCrArCrGrUrArGrArArCrArGrArUrGrCrArUrGrArCrGrCrCrA-5’), SK2 (hs.Ri.SPHK213.1; duplex sequences:

5’-rCrCrCrUrGrArArArCrUrArArArCrArArGrCrUrUrGrGrUAC-3’,

3’-rGrUrArCrCrArArGrCrUrUrGrUrUrUrArGrUrUrUrCrArGrGrGrCrU-5’), and negative control dsiRNA (51-01-14-04) were purchased from Integrated DNA Technologies, Inc. (Coralville, Iowa).

### Viral pseudotype and viral-like particle production

Murine leukemia virus and lentivirus pseudotypes were prepared by co-transfecting HEK293T cells with the packaging plasmid gag-pol or psPAX2, MLV retroviral or lentiviral vector encoding LacZ, and a plasmid encoding the glycoprotein of interest (EBOV Δmucin GP, MARV GP, LFV GP, NiV F and G (1:1), Junin GPC, IAV envelope proteins (Vict HA, 1918 NA), SARS-CoV-1 Spike, MERS Spike, or VSV-G) at a 1:1:1.25 ratio respectively. Similarly, EBOV viral-like particles (VLPs) were prepared by co-transfecting HEK293T cells with plasmids encoding the EBOV nucleoprotein (NP), EBOV VP40 fused to β-lactamase (βlam) or GFP, and the viral glycoprotein of interest (EBOV Δmucin GP, EBOV Δmucin GP^F535R^, or VSV G) at a 1:1:1.25 ratio. Transfections were performed using the jetPRIME transfection reagent (Polyplus transfection) according to the manufacturers protocol. Cell supernatants were harvested at 48, 72, and 96 h post-transfection. MLVs and lentiviruses supernatants were passed through a 0.45µm filter and VLPs supernatants pre-cleared by centrifugation at 4,000g for 5min. Pseudotypes and VLPs were then concentrated by ultracentrifugation (20,000 RPM, 4 ºC, 1.5h, Beckman Coulter Optima XPN-100, SW32Ti rotor) through a 20% (*w/v*) sucrose cushion. Pellets were re-suspended in PBS, aliquoted, and stored at -80 ºC.

### Virus entry assays

For MLV and lentivirus pseudotype transduction assays, HT1080 cells were seeded and grown to approximately 60% confluency in white 96-well pates. Cells were pre-incubated with inhibitor or vehicle (DMSO) in serum-free MEM containing 5 µg/mL polybrene. MLV or lentivirus pseudotypes encoding LacZ and harbouring the viral fusion protein of interest were added to the media. 4-6 hours after virus addition, the media was replaced with MEM containing 15 mM NH_4_Cl and supplemented with 10% FBS (Sigma), 0.3 mg/mL L-glutamine, 100 U/mL penicillin, and 100µg/mL streptomycin (Wisent). Approximately 12 hours later, the media was replaced with complete MEM (without NH_4_Cl) and cells incubated an additional 48 hours Transduced (LacZ+) cells were quantified using the Beta-Glo Assay System (Promega, Madison, Wisconsin) following the manufacturers’ protocol. Luminescence was measured using a Synergy Neo2 multi-mode plate reader (BioTek Instruments, Winooski, Vermont).

For the VLP entry assays, HT1080 cells were seeded and grown to 90% confluency. For experiments with inhibitors, cells were pre-treated with the inhibitors or vehicle (DMSO) for 1 hour in serum-free MEM. For S1P experiments, cells were pre-treated for 1 hour in serum-free MEM with inhibitor or vehicle, and S1P (125 nM) was added 10 minutes prior to virus addition. VP40-βlam VLPs harbouring EBOVΔM or VSV-G were added at a MOI between 0.2 and 0.4. 3 hours post infection, cells were loaded with a β-lactamase cleavable FRET substrate, CCF2-AM (ThermoFisher Scientific, Waltham, Massachusetts), according to manufacturer’s protocol and supplemented with 15 mM NH_4_Cl and 250 µM probenecid (Sigma). Cells were then incubated for 1h at room temperature in the dark, washed with PBS, trypsinized, and resuspended with 2% FBS in PBS prior to analysis by flow cytometry (FACSCelesta or LSRFortessa, BD Biosciences). Flow cytometry analysis was performed using the FlowJo software (BD Biosciences) and infection was quantified by using uninfected controls to assess the percentage of cells that underwent a shift from 530 nm to 460 nm emission, representing cleaved CCF2.

### Sphingosine kinase 1/2 activity assay

SK1 and SK2 specific master mixes were pre-equilibrated at room temperature and prepared immediately before use. SK1 reaction buffer contained 100nM SK1 (R&D systems, Minneapolis, Minnesota), 25uM NBD-Sph (Cayman Chemical Company, Ann Arbor, Michigan), 30 mM Tris-HCl, pH 7.4, 0.05% Triton X-100, 150 mM NaCl, 10% glycerol, 1 mM Na3VO4, 10 mM NaF, and 10 mM b-glycero-phosphate and SK2 reaction buffer contained 15nM SK2 (R& D systems, Minneapolis, Minnesota), 75 NBD-Sph, 30 mM Tris-HCl, pH 7.4, 0.05% Triton X-100, 200 mM KCl, and 10% glycerol. Solutions were vortexed and 18 ul of the master mix was added per well of a 384-well black plates (ThermoFisher Scientific, Waltham, Massachusetts), followed by the addition of 1ul of inhibitors at 20uM, for a final concentration of 1uM. The assay was initiated with 1ul of 20× ATP-Mg (20 mM ATP, 200 mM MgCl2, 900 mM Tris-HCl, pH 7.4), after which the 384-well plate was shaken at 800 rpm for 1 min to allow for complete mixing. After an incubation of 10 min (SK2) and 30 min (SK1) at room temperature, fluorescence emission was measured with a Synergy Neo2 multi-mode plate reader (BioTek Instruments, Winooski, Vermont) where the excitation wavelength was set at 550 nm, and emission wavelength at 584 nm.

### Pre-cleaved virus

βlam VLPs harboring EBOV Δmucin GP were incubated in 0.2 mg/mL thermolysin (MilliporeSigma, Burlington, Massachusetts) or PBS (mock) for 30 minutes at 37ºC. Phosphoramidon (MilliporeSigma, Burlington, Massachusetts) was then added to a final concentration of 500 µM and the mixture was incubated on ice for 10 minutes. Pre-cleaved (thermolysin-treated) and mock-cleaved (PBS-treated) samples were aliquoted and stored at -80 ºC.

### DsiRNAs transfection and infection assays

HT1080 were cells grown to 90% confluency and transfected with the DsiRNAs using Lipofectamine RNAiMAX (ThermoFisher Scientific, Waltham, Massachusetts) according to the manufacturer’s protocol. 24 hours post-transfection, cells were re-seeded for VLP entry assays, which were conducted 48 hours post-transfection as described above, with the exception that cells were pre-incubated in serum-free MEM that did not contain any inhibitor or vehicle. In addition, 48 hours post-transfection, cells seeded for lysate preparation were washed once with PBS, followed by the addition of lysis buffer (1% Triton X-100, 0.1% IGEPAL CA-630, 150mM NaCl, 50mM Tris-HCl, pH 7.5) containing protease inhibitors (Cell Signaling Technology, Danvers, Massachusetts). Proteins were resolved on SDS-polyacrylamide gels and transferred to polyvinylidenedifluoride (PVDF) membranes. Membranes were blocked for 1h at room temperature with blocking buffer (5% skim milk powder dissolved in 25mM Tris, pH 7.5, 150mM NaCl, and 0.1% Tween-20 [TBST]). PVDF membranes were then incubated overnight at 4 ºC with the appropriate primary antibody. The following day, blots were washed in TBST and incubated with HRP-conjugated secondary antibody for 1h at room temperature. PVDF membranes were then washed again, incubated in chemiluminescence substrate, and imaged using the ChemiDoc XRS+ imaging system (Bio-Rad Laboratories, Hercules, California).

### Replicative EBOV and SARS-CoV-2 Infection assays

For infection assays with replication-competent EBOV, Vero cells were seeded onto black 96 well plates with glass bottom wells. After 48 hours, cells were pre-treated for 1 hour with vehicle (DMSO) or PF-543 in DMEM supplemented with 1% FBS. Ebola virus (Mayinga) expressing GFP was added to media containing inhibitors or vehicle and incubated together for 15 minutes prior to addition of the mixture to the cells at a final MOI of 1. Control wells containing no virus were also included. After infection, GFP expression was monitored daily using a BioTek Synergy/HTX plate reader (BioTek Instruments, Winooski, Vermont). These experiments were performed at the CL-4 facility at the National Microbiology Laboratory of the Public Health Agency of Canada.

For infection assays with replication-competent SARS-CoV-2 (hCoV-19/Canada/ON-VIDO-01/2020, GISAID accession# EPI_ISL_425177), Huh7.5 cells were seeded and once adhered, inhibitor or vehicle was incubated with the cells at 2x concentration for 1 hour (71). SARS-CoV-2 was then added to each well at an MOI of 0.01 and to give a final inhibitor concentration of 1x. The inoculum + drug mixture was incubated on cells for 2-3 hours at 37 ºC, cells were then washed with MEM and media containing inhibitor or vehicle was added back to cells and incubated for an additional 48h at 37 ºC. Supernatant was then collected and used directly to measure the TCID50/mL on Vero cells. These experiments were performed at the CL-4 facility at the National Microbiology Laboratory of the Public Health Agency of Canada.

### S1PR downstream signaling assay

HT1080 cells were seeded and grown to 50% confluency, washed with PBS, and pre-treated for 1 hour with inhibitor or vehicle (DMSO) in serum-free MEM. Cells were then stimulated for 30 minutes with S1P prepared in 4 mg/mL fatty acid free Bovine Serum Albumin (BSA) (MilliporeSigma) in PBS or mock-treated for 30 minutes with 4 mg/mL fatty acid free BSA in PBS. Cells were then washed once with cold PBS and lysed in cold lysis buffer (1% Triton X-100, 0.1% IGEPAL CA-630, 150mM NaCl, 50mM Tris-HCl, pH 7.5) containing protease and phosphatase inhibitors (Cell Signaling). Proteins were resolved on SDS-polyacrylamide gels (Bio-Rad) and transferred to polyvinylidenedifluoride (PVDF) membranes. Membranes were blocked at RT with blocking buffer (5% skim milk powder in TBST containing sodium orthovanadate (Na_3_VO_4_, 1mM, Alfa Aesar) and sodium fluoride (NaF, 10mM, VWR)). After 1 hour of blocking, membranes were incubated overnight at 4ºC with either the pAkt antibody in 5% BSA in TBST containing Na_3_VO_4_ and NaF, or Akt antibody in 5% skim milk powder in TBST. The following day, blots were washed in TBST and incubated with HRP-conjugated secondary antibody for 1h at room temperature. After 3 more washes in TBST, the membranes were incubated in chemiluminescence substrate and imaged using the ChemiDoc XRS+ imaging system (Bio-Rad Laboratories).

### Attachment and internalization assays

For the attachment assay, HT1080 cells were seeded and grown to 90% confluency. Cells were pre-treated with serum-free MEM containing inhibitor or vehicle for 1 hour at 37 ºC. The cells were detached with 5 mM EDTA and resuspended in 2% FBS in PBS containing inhibitor or vehicle. After incubation at 4 ºC for 15 minutes, VP40-GFP EBOVΔM VLPs were added on ice and incubated for 1 h at 4 ºC to allow the virus to attach to the cell surface. As a control, one set of vehicle treated cells was incubated with trypsin to remove bound virions. The cell and virus mixtures were spun down and washed three times with cold PBS prior to resuspension in 2% FBS in PBS containing SYTOX Red dead cell stain (ThermoFisher) and analysis by flow cytometry.

For the internalization experiments, HT1080 cells were seeded and grown to 90% confluency. Cells were pre-treated with serum-free MEM containing inhibitor or vehicle (DMSO) for 1 hour at 37ºC. The cells were then incubated at 4ºC for 15 minutes prior to spinoculation of VP40-GFP EBOVΔM VLPs at 300 x g for 30 minutes. Cells were then washed 3x with cold PBS to removed unbound virions, pre-warmed media containing inhibitor or vehicle was added, and cells were moved to 37 ºC for 1 hour to allow for internalization. Cells were then washed again with cold PBS and incubated with 0.5% trypsin-EDTA (Gibco) at 4 ºC for 30 minutes to remove VLPs that were still at the surface. Cells were then distributed into tubes containing cold PBS, spun down and washed twice with cold PBS, and finally resuspended in 2% FBS in PBS containing SYTOX Red dead cell stain (ThermoFisher) prior to analysis by flow cytometry.

### Fluorescence microscopy and image analysis

HT1080 cells were seeded onto coverslips that were coated with Poly-D-lysine (Sigma) and grown to approximately 50% confluency. Cells were then pre-treated with inhibitors or vehicle (DMSO) in serum-free MEM for 1 h followed by addition of VP40-GFP VLPs harboring the fusion deficient EBOV Δmucin GP^F535R^. Cells were incubated for 2.5 hours at 37 ºC, CellTracker Blue CMAC dye (ThermoFisher) was added according to manufacturer’s protocol, and cells were incubated for an additional 30 minutes. Cells were then washed with PBS, fixed with formalin, permeabilized with 0.5% triton X-100, and blocked with 20% FBS in PBS for 30 minutes. Cells were then incubated with a NPC1 primary antibody (1:70) followed by a DY650 secondary antibody (1:400) and mounted with PermaFluor Aqueous Mounting medium (ThermoFisher). Imaging was performed with a LSM800 confocal microscope (AxioObserver Z1, Zeiss) using a 63x / 1.4NA oil Plan Apochromat objective. Approximately fifteen z-stacks were acquired per image with a pixel size of 0.1 µm.

Image analysis was performed using Imaris software v. 8.4.2 (Bitplane). In brief, each cell was modeled based on the CMAC cytoplasmic stain using the surfaces module, and VLPs were modeled as spots. The number of spots per cell was then determined and the modeled spots were assigned colocalization values based on intensity correlation to NPC1. Intensity thresholds were manually set for each experiment but kept constant between experimental conditions. By dividing the number of spots above the colocalization threshold by the total number of spots per cell, the percentage of VLPs colocalizing with NPC1 was determined.

## Supporting information

Supporting Information: Compound Synthesis

## Acknowledgments

We would like to thank the uOttawa CBIA core staff, Dr. Chloë van Oostende, Skye Green, and Redaet Daniel, as well as Vera Tang from the uOttawa Flow Cytometry and Virometry core for their technical support. We would also like to thank and acknowledge Dr. Shirley Qiu for technical advice and contribution to editing the manuscript and Dr. Suresh Gadde for providing advice and help with the manuscript.

This research was funded by the Canadian Institutes of Health Research: grant number ER1-143489352509 and PJT155984390487 to M.C. and D.K., grant number 143521 to G.P.K., and part of this research was also supported by a COVID-19 Rapid Research grant (CIHR, OV3 170632) to M.C. and D.K.; C.M.S. was supported by a graduate scholarship from the Natural Sciences and Engineering Research Council of Canada and an Ontario Graduate Scholarship. K. Y-P A. was supported by a Centre for Catalysis Research Innovation (CCRI) Collaborative Research Award for Medicinal Chemistry. Support from the University of Ottawa to A. M. B. is gratefully acknowledged. K.F was supported by an Ontario Graduate Scholarship. M.C. is a Canada Research Chair in Molecular Virology and Antiviral Therapeutics (950-232840) and recipients of Ontario Ministry of Research, Innovation and Science Early Researcher Awards (ER18-14-091). Additional support is provided to D.K. by the Public Health Agency of Canada.

