## Supporting Information: Compound Synthesis for "Sphingosine kinases promote Ebola virus infection and can be targeted to inhibit filoviruses, coronaviruses, and arenaviruses using late endocytic trafficking to enter cells"

***Synthesis of PF-543 derivatives***

### Overall synthesis

The overall synthetic scheme is shown below

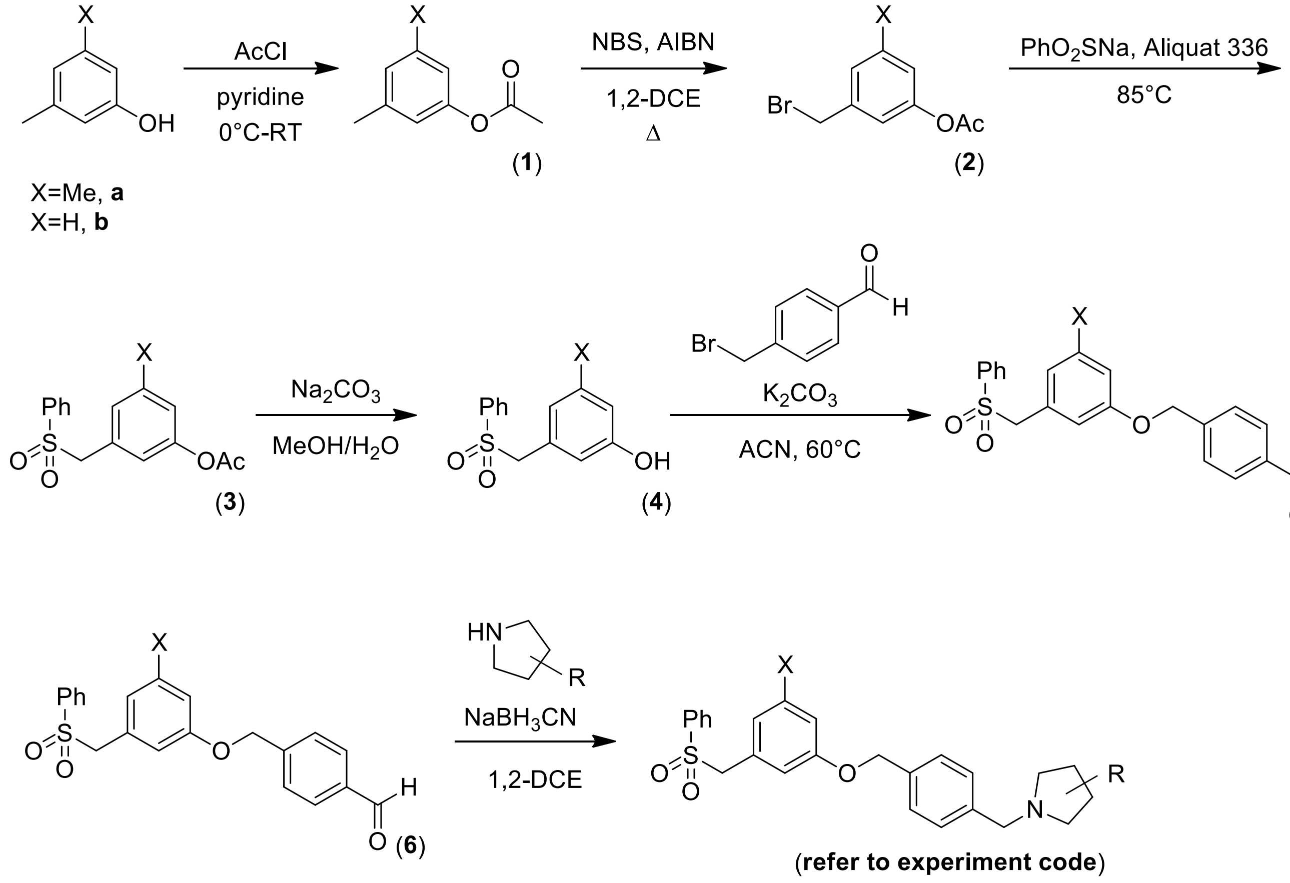


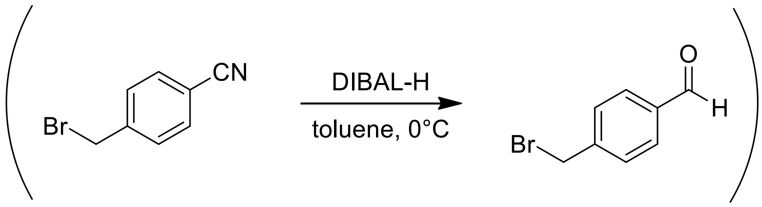

**Synthesis**

The majority of the synthesis is adapted from Bioorg. Med. Chem. 2014 Vogt, unless indicated otherwise.

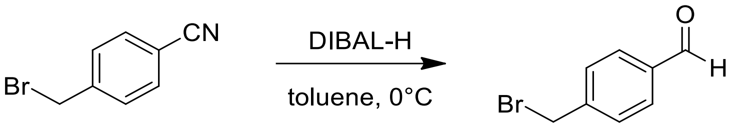


To a solution of α-bromo-*p*-tolunitrile (1.0 eq) in toluene (0.5 M), cooled to 0°C and placed under argon, was added DIBAL-H (1.0 in toluene, 1.4 eq) prior to stirring at 0°C for one hour. The reaction was quenched by the addition of chloroform and 10% aqueous hydrochloric acid and stirring was maintained at 0°C for a second hour. The organic phase was isolated and washed with water prior to being dried over anhydrous magnesium sulfate, filtered and evaporated to dryness to obtain a white powder in an average **yield** of 80%. $\mathbf{R_f}$ (9:1 hex:EtOAc) ~0.28. **$^1$H NMR** (400 MHz, $CDCl_3$) δ 10.03 (s, 1H), 7.88 (d, *J*=8.2 Hz, 2H), 7.57 (d, *J*=8.1 Hz, 2H), 4.53 (s, 2H). **$^{13}$C NMR** (100 MHz, $CDCl_3$) δ 191.5, 144.2, 136.1, 130.2, 129.7, 31.9. **HRMS (EI)** calc. for [$C_8H_7OBr$] 197.9680 Da, obt. 197.9687 Da.
(*Mol. Pharmaceutics* 2013 Zhang)

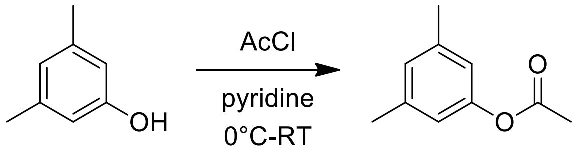


(**1a**)

To a solution of the phenol (1.0 eq) in pyridine (1.67 M), cooled to 0°C, was added dropwise the acid chloride (1.2 eq), resulting in the formation of a pale precipitate. The resulting heterogeneous mixture was stirred for 18 hours as it gradually warmed to room temperature and formed a dark orange solution. This solution was poured onto water prior to extracting the product with ethyl acetate. The combined organic extracts were dried over anhydrous magnesium sulfate, filtered, and evaporated under reduced pressure prior to being purified by flash column chromatography (100:0 to 9:1 hexanes:ethyl acetate). The average **yield** was 90% as a clear colourless oil. $\mathbf{R_f}$ (9:1) ~0.39. **$^1$H NMR** (400 MHz, $CDCl_3$) δ 6.88 (s, 1H), 6.71 (s, 2H), 2.32 (s, 6H), 2.29 (s, 3H). **$^{13}$C NMR** (100 MHz, $CDCl_3$) δ 169.7, 150.6, 139.3, 127.6, 119.1, 21.2, 21.1.

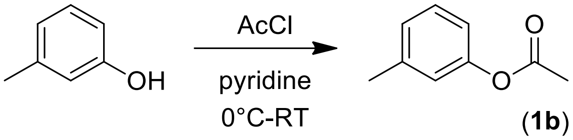


(**1b**)

Compound **1b** was prepared in the same manner as **1a** above, and purified by flash column chromatography (9:1 hexanes:ethyl acetate). The **yield** was 89% as a clear colourless oil. $\mathbf{R_f}$ (9:1) ~0.43. $^{1}$**H NMR** (400 MHz, $CDCl_3$) δ 7.29-7.25 (m, 1H), 7.07-7.04 (m, 1H), 6.92-6.88 (m, 2H), 2.37 (s, 3H), 2.30 (s, 3H). $^{13}$**C NMR** (100 MHz, $CDCl_3$) δ 169.6, 150.6, 139.6, 129.1, 126.6, 122.1, 118.5, 21.3, 21.1.

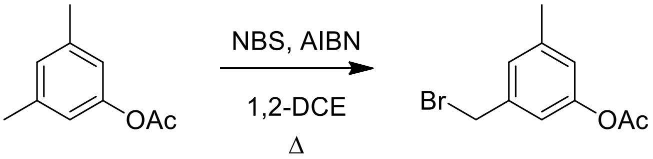


(**2a**)

To a solution of the phenyl acetate (1.0 eq) in 1,2-dichloroethane (1.0 M) were added *N*-bromosuccinimide (0.8 eq) and azobisisobutyronitrile (17.6 meq) prior to heating to reflux (83°C) and stirring for one hour. Upon cooling to room temperature, a white precipitate crashed out; it was removed by filtration and rinsed with diethyl ether. The filtrate and rinses were combined and evaporated under reduced pressure prior to being purified by flash column chromatography (190:10 to 9:1 hexanes:ethyl acetate). The average **yield** was 71% as a clear colourless oil. $\mathbf{R_f}$ (9:1) ~0.39. $^{1}$**H NMR** (400 MHz, $CDCl_3$) δ 7.08 (s, 1H), 6.95 (s, 1H), 6.86 (s, 1H), 4.44 (s, 2H), 2.36 (s, 3H), 2.30 (s, 3H). $^{13}$**C NMR** (100 MHz, $CDCl_3$) δ 169.4, 150.7, 140.1, 138.9, 127.3, 122.3, 119.3, 32.7, 21.2, 21.1.

(Vogt and *J. Org. Chem.* 1977 Newman)

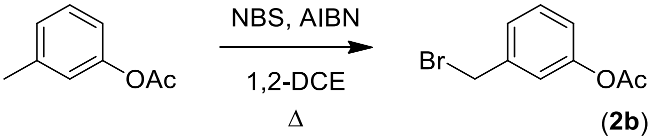


(**2b**)

Compound **2b** was prepared in the same manner as **2a** above, and purified by flash column chromatography (95:5 hexanes:ethyl acetate). The **yield** was 43% as a pale yellow oil. $\mathbf{R_f}$ (95:5) ~0.16. **$^{1}$H NMR** (400 MHz, $CDCl_3$) δ 7.36 (dd, *J*=7.9 Hz, *J*=7.8 Hz, 1H), 7.28-7.26 (m, 1H), 7.15 (dd, *J*=2.0 Hz, *J*=1.9 Hz, 1H), 7.06-7.03 (m, 1H), 4.48 (s, 2H), 2.31 (s, 3H). **$^{13}$C NMR** (100 MHz, $CDCl_3$) δ 169.3, 150.7, 139.3, 129.8, 126.4, 122.2, 121.6, 32.5, 21.1.

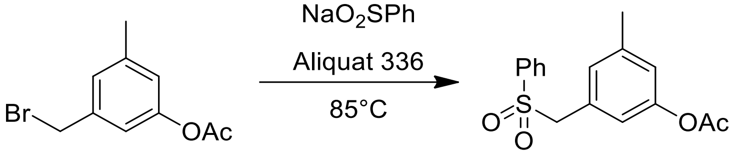

(**3a**)

A slurry of the benzyl bromide (1.0 eq), benzenesulfinic acid sodium salt (1.1 eq) and Aliquat 336 (20 meq) was heated to 85°C and stirred for 28 hours. Upon cooling to room temperature, the product was resuspended in ethyl acetate, using sonication, and filtered over Celite. This solution was evaporated under reduced pressure prior to being purified by flash column chromatography (0-7% ethyl acetate in DCM). The average **yield** was 72% as a clear colourless oil. $\mathbf{R_f}$ (95:5 DCM:ethyl acetate) ~0.55. **$^{1}$H NMR** (400 MHz, $CDCl_3$) δ 7.69-7.66 (m, 2H), 7.64-7.60 (m, 1), 7.50-7.46 (m, 2H), 6.87 (s, 1H), 6.76 (s, 1H), 6.66 (s, 1H), 4.26 (s, 2H), 2.27 (s, 3H), 2.25 (s, 3H). **$^{13}$C NMR** (100 MHz, $CDCl_3$) δ 169.2, 150.5, 139.8, 137.7, 133.7, 129.2, 129.1, 128.9, 128.6, 122.7, 121.0, 62.4, 21.1, 21.0.
(Vogt and *Biochem. J.* 2012 Schnute)

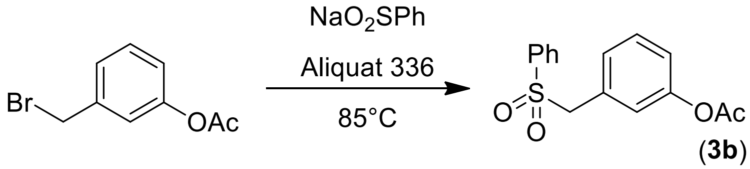

(**3b**)

Compound **3b** was prepared in the same manner as **3a** above, and purified by flash column chromatography (100% DCM to 9:1 DCM:ethyl acetate). The **yield** was 63% as a white powder. $\mathbf{R_f}$ (100% DCM) ~0.26. **$^{1}$H NMR** (400 MHz, $CDCl_3$) δ 7.67-7.60 (m, 3H), 7.49-7.46 (m, 2H), 7.27 (t, overlaps $CHCl_3$, 1H), 7.08-7.05 (m, 1H), 6.94-6.90 (m, 2H), 4.31 (s, 2H), 2.28 (s, 3H). **$^{13}$C NMR** (100 MHz, $CDCl_3$) δ 169.1, 150.7, 137.7, 133.8, 129.7, 129.5, 129.0, 128.6, 128.2, 124.0, 122.1, 62.4, 21.0.

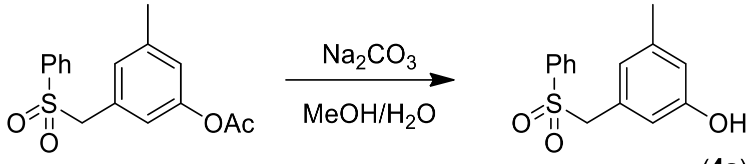


(**4a**)

To a solution of the phenyl acetate (1.0 eq) in methanol (0.5 M) and water (0.25 M) was added saturated aqueous sodium bicarbonate (0.25 M) prior to stirring at room temperature for 21 hours, at which point TLC analysis showed complete reactant consumption. The organic solvent was removed by evaporation under reduced pressure and the aqueous phase was acidified to pH 1 using 6N HCl. The product was extracted with ethyl acetate and the combined organic extracts were washed with water and brine prior to being dried over anhydrous magnesium sulfate, filtered, and evaporated to dryness. The average **yield** was 86% as an off-white powder. **$^1$H NMR** (400 MHz, $CDCl_3$) δ 7.70-7.68 (m, 2H), 7.63-7.60 (m, 1H), 7.50-7.46 (m, 2H), 6.63 (s, 1H), 6.48 (s, 1H), 6.38 (s, 1H), 4.23 (s, 2H), 2.18 (s, 3H). **$^{13}$C NMR** (100 MHz, $CDCl_3$) δ 155.8, 139.9, 137.8, 133.8, 128.9, 128.6, 124.0, 116.7, 114.8, 62.7, 14.1.

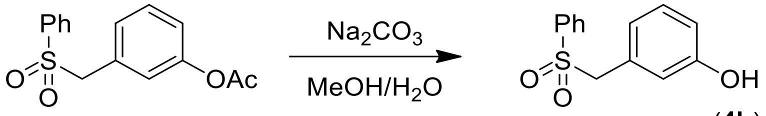


(**4b**)

Compound **4b** was prepared in the same manner as **4a** above, and obtained in 92% **yield** as a white powder. **$^1$H NMR** (400 MHz, $CDCl_3$) δ 7.67 (dd, not fully resolved, 2H), 7.62 (dd, *J*=7.4 Hz, *J*=7.4 Hz, 1H), 7.48 (dd, *J*=8.0 Hz, *J*=7.5 Hz, 2H), 7.11 (dd, *J*=7.8 Hz, *J*=7.8 Hz, 1H), 6.81 (dd, *J*=8.0 Hz, *J*=2.4 Hz, 1H), 6.68 (dd, *J*=2.0 Hz, *J*=1.9 Hz, 1H), 6.58 (d, *J*=7.6 Hz, 1H), 4.27 (s, 2H). **$^{13}$C NMR** (100 MHz, $CDCl_3$) δ 155.7, 137.8, 133.8, 129.8, 129.5, 128.9, 128.6, 123.2, 117.6, 116.0, 62.7.

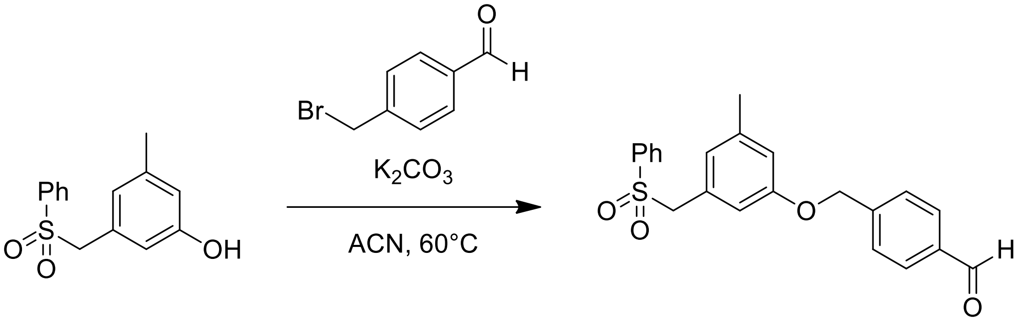


(**5a**)

To a solution of the phenol (1.0 eq) and benzyl bromide (1.0 eq) in acetonitrile (0.2 M) was added potassium carbonate (5.0 eq) prior to heating the reaction mixture to 60°C and stirring for 2 hours. Following cooling to room temperature, the reaction mixture was diluted with ethyl acetate and 50% aqueous brine. The organic phase was additionally washed with brine prior to being dried over anhydrous magnesium sulfate, filtered, and evaporated under reduced pressure. The product was purified by flash column chromatography (100% DCM to 9:1 DCM:ethyl acetate). The average **yield** was 56% as an off-white powder. **R$_f$** (95:5 DCM:ethyl acetate) ~0.73. **$^1$H NMR** (400 MHz, $CDCl_3$) δ 10.04 (s, 1H), 7.92 (d, *J*=8.0 Hz, 2H), 7.68 (d, *J*=7.9 Hz, 2H), 7.62 (dd, *J*=7.5 Hz, *J*=7.4 Hz, 1H), 7.57 (d, *J*=7.9 Hz, 2H), 7.48 (dd, *J*=7.8 Hz, *J*=7.8 Hz, 2H), 6.76 (s, 1H), 6.56 (s, 1H), 6.50 (s, 1H), 5.05 (s, 2H), 4.24 (s, 2H), 2.24 (s, 3H). **$^{13}$C NMR** (100 MHz, $CDCl_3$) δ 191.8, 158.3, 143.7, 139.9, 138.1, 136.0, 133.7, 130.0, 129.2, 128.9, 128.7, 127.4, 124.8, 116.4, 113.9, 69.1, 62.8, 21.3.

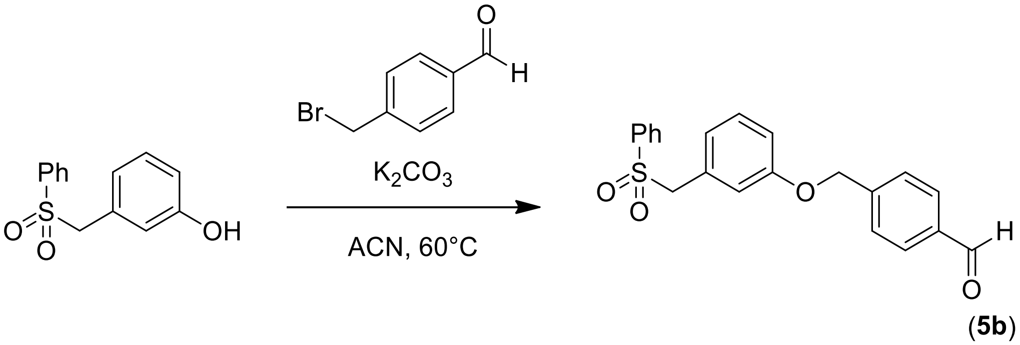


(**5b**)

Compound **5b** was prepared in the same manner as **5a** above, and purified by flash column chromatography (100% DCM to 9:1 DCM:ethyl acetate). The product was obtained in 66% **yield** as a white powder. **R$_f$** (100% DCM) ~0.14. **$^1$H NMR** (400 MHz, $CDCl_3$) δ 10.04 (s, 1H), 7.92 (d, *J*=8.2 Hz, 2H), 7.67-7.57 (m, 5H), 7.49-7.45 (m, 2H), 7.17 (dd, *J*=7.9 Hz, *J*=7.9 Hz, 1H), 6.93

(ddd, $J$=8.3 Hz, $J$=2.5 Hz, $J$=0.78 Hz, 1H), 6.80 (dd, $J$=2.2 Hz, $J$=1.8 Hz, 1H), 6.66 (d, $J$=7.6 Hz, 1H), 5.08 (s, 2H), 4.29 (s, 2H). **$^{13}$C NMR** (100 MHz, $CDCl_3$) δ 191.8, 158.3, 143.6, 137.9, 136.0, 133.7, 130.0, 129.7, 129.6, 128.9, 128.6, 127.5, 123.8, 116.9, 115.6, 69.2, 62.8.

**Final compounds**

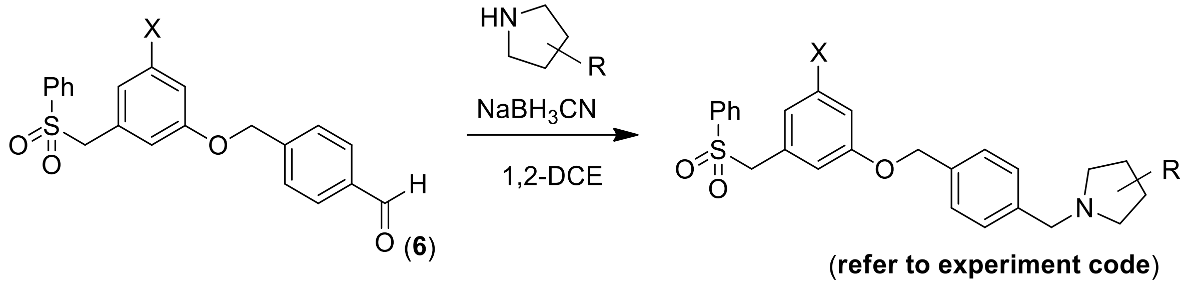


(**refer to experiment code**)

General procedure: To a solution of the pyrrolidine (1.1 eq) in 1,2-dichoroethane (0.2 M) was added the aldehyde (**5a** or **5b**, 1.0 eq) prior to stirring at room temperature for one hour, after which sodium cyanoborohydride (1.5 eq) was added and stirring was continued at room temperature for 16 hours. The reaction mixture was diluted with DCM and washed with saturated aqueous sodium bicarbonate. The organic phase was collected and evaporated under a constant stream of compressed air prior to purifying the product by flash column chromatography. Some reactions were catalysed using a drop of glacial acetic acid; these will be indicated individually.

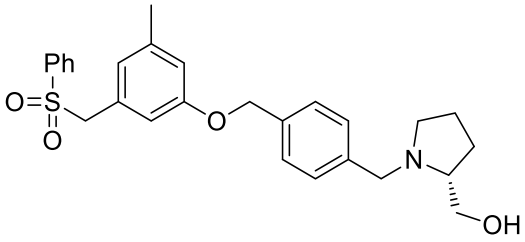


**PF-543**
(**AMB-KA-016**)

Compound **KA-016 (PF-543)** was prepared from **5a** and (*R*)-(-)-2-prolinol, purified using a 0-10% gradient of methanol in DCM, and obtained in 17% **yield** as a yellow oil. **$R_f$** (9:1 DCM:MeOH) ~0.27. **$^{1}$H NMR** (400 MHz, $CDCl_3$) δ 7.67 (dd, $J$=8.5 Hz, $J$=1.4 Hz, 2H), 7.65-7.61 (m, 1H), 7.50-7.44 (m, 4H), 7.40 (d, $J$=8.0 Hz, 2H), 6.76 (s, 1H), 6.52 (s, 1H), 6.48 (s, 1H), 4.94 (s, 2H), 4.23 (s, 2H), 4.18 (d, $J$=13.2 Hz, 1H), 3.77-3.71 (m, 2H), 3.63 (dd, $J$=12.0 Hz,

$J$=4.3 Hz, 1H), 3.23 (br s, 1H), 3.10 (br s, 1H), 2.58 (q, $J$=8.1 Hz, 1H), 2.23 (s, 3H), 2.03-1.80 (m, 4H). **$^{13}$C NMR** (100 MHz, $CDCl_3$) δ 158.6, 139.8, 138.0, 133.7, 130.0, 129.0, 128.9, 128.7, 127.7, 124.6, 116.5, 113.8, 69.5, 66.3, 62.9, 61.4, 58.6, 54.1, 27.1, 23.5, 21.3. **HRMS (ESI$^+$)** calc. for $[C_{27}H_{31}NO_4S+H]^+$ 466.2052, obt. 466.2044.

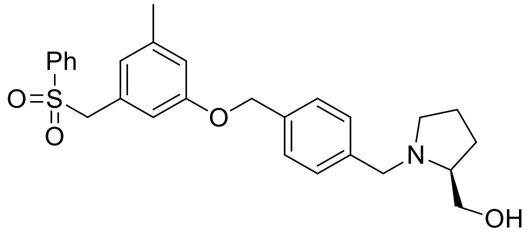


(**AMB-KA-018**)

Compound **KA-018 (Compound 18)** was prepared from **5a** and (*S*)-pyrrolidin-2-ylmethanol, purified using a 0-10% gradient of methanol in DCM, and obtained in 12% **yield** as a clear oil. **$R_f$** (9:1 DCM:MeOH) ~0.47. **$^1$H NMR** (400 MHz, $CDCl_3$) δ 7.67 (dd, ill-defined, 2H), 7.65-7.61 (m, 1H), 7.50-7.46 (m, 2H), 7.43-7.37 (m, 4H), 6.75 (s, 1H), 6.52 (s, 1H), 6.48 (s, 1H), 4.93 (s, 2H), 4.24 (s, 2H), 4.13 (d, $J$=13.0 Hz, 1H), 3.74-3.56 (m, 3H), 3.18-3.14 (m, 1H), 3.00 (br s, 1H), 2.50 (q, $J$=12.9 Hz, 1H), 2.23 (s, 3H), 2.00-1.77 (m, 4H). **$^{13}$C NMR** (100 MHz, $CDCl_3$) δ 158.6, 139.8, 138.0, 136.5, 133.6, 129.7, 129.0, 128.8, 128.7, 127.7, 124.5, 116.5, 113.8, 69.6, 65.6, 62.9, 61.5, 58.4, 54.2, 27.3, 23.4, 21.3. **HRMS (ESI$^+$)** calc. for $[C_{27}H_{31}NO_4S+H]^+$ 466.2052, obt. 466.2066.

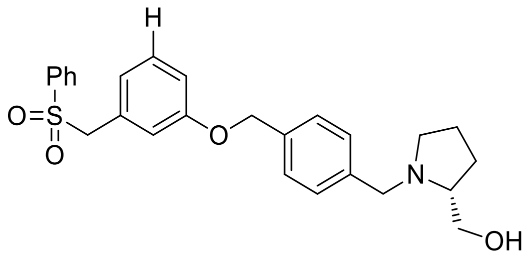


(**AMB-KA-019**)

Compound **KA-019 (Compound 19)** was prepared from **5b** and (*R*)-(-)-2-prolinol, purified using a 0-10% gradient of methanol in DCM and obtained in 19% **yield** as a yellow oil. **$R_f$** (9:1 DCM:MeOH) ~0.44. **$^1$H NMR** (400 MHz, $CDCl_3$) δ 7.67-7.60 (m, 3H), 7.47 (dd, $J$=8.0 Hz, $J$=7.6 Hz, 2H), 7.40 (q, $J$=8.2 Hz, $J$=7.4Hz, 4H), 7.16 (dd, $J$=8.0 Hz, $J$=7.8 Hz, 1H), 6.94-6.91

(m, 1H), 6.75 (dd, $J$=2.2 Hz, $J$=1.8 Hz, 1H), 6.64 (br d, $J$=7.6 Hz, 1H), 4.96 (s, 2H), 4.28 (s, 2H), 4.14 (d, $J$=13.1 Hz, 1H), 3.72 (dd, $J$=11.6 Hz, $J$=3.3 Hz, 1H), 3.63 (d, $J$=13.1 Hz, 1H), 3.58 (dd, $J$=11.6 Hz, $J$=3.8 Hz, 1H), 3.17-3.13 (m, 1H), 2.53-2.47 (m, 1H), 2.01-1.77 (m, 4H). **$^{13}$C NMR** (100 MHz, $CDCl_3$) δ 158.6, 137.9, 136.4, 133.7, 129.7, 129.6, 129.4, 128.9, 128.6, 127.7, 123.5, 116.9, 115.7, 69.6, 65.6, 62.8, 61.5, 58.4, 54.2, 27.3, 23.4. **HRMS (ESI$^+$)** calc. for $[C_{26}H_{29}NO_4S+H]^+$ 452.1896, obt. 452.1896.

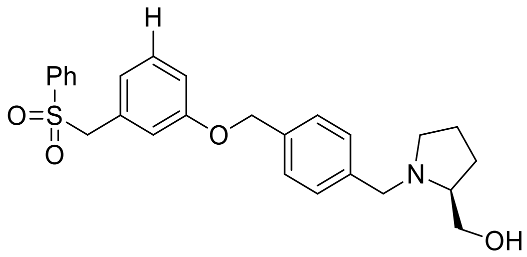


(**AMB-KA-020**)

Compound **KA-020 (Compound 20)** was prepared from **5b** and (*S*)-pyrrolidin-2-ylmethanol, purified using a 0-10% gradient of methanol in DCM, and obtained in 17% **yield** as a yellow oil. **$R_f$** (9:1 DCM:MeOH) ~0.35. **$^1$H NMR** (400 MHz, $CDCl_3$) δ 7.65 (dd, ill-defined, 2H), 7.63-7.60 (m, 1H), 7.48 (d, $J$=8.0 Hz, 2H), 7.46-7.44 (m, 2H), 7.40 (d, $J$=8.1 Hz, 2H), 7.16 (dd, $J$=8.0 Hz, $J$=7.8 Hz, 1H), 6.92 (dd, $J$=8.3 Hz $J$=1.8 Hz, 1H), 6.75 (dd, $J$=2.2 Hz, $J$=1.9 Hz, 1H), 6.64 (d, $J$=7.5 Hz, 1H), 4.94 (s, 2H), 4.28 (s, 2H), 4.15 (d, $J$=13.0 Hz, 1H), 3.73 (dd, $J$=11.7 Hz, $J$=3.2 Hz, 1H), 3.67 (d, $J$=11.8 Hz, 1H), 3.60 (dd, $J$=11.6 Hz, $J$=3.6 Hz, 1H), 3.19 (br s, 1H), 3.04 (br s, 1H), 2.56-2.48 (m, 1H), 2.02-1.78 (m, 4H). **$^{13}$C NMR** (100 MHz, $CDCl_3$) δ 158.6, 137.9, 133.7, 129.8, 129.6, 129.4, 128.9, 128.6, 127.7, 123.5, 116.9, 115.7, 69.6, 62.8, 61.4, 58.5, 54.2, 27.2, 23.5. **HRMS (ESI$^+$)** calc. for $[C_{26}H_{29}NO_4S+H]^+$ 452.1896, obt. 452.1899.

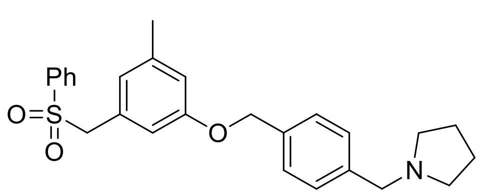


(**AMB-KA-035**)

Compound **KA-035 (Compound 35)** was prepared from **5a** and pyrrolidine, purified using a 0-10% gradient of methanol in DCM, and obtained in 12% **yield**. **$R_f$** (9:1 DCM:MeOH) ~0.62. **$^1$H**

**NMR** (400 MHz, $CDCl_3$) δ 7.68-7.62 (m, 3H), 7.51-7.42 (m, 6H), 6.76 (s, 1H), 6.53 (s, 1H), 6.47 (s, 1H), 4.96 (s, 2H), 4.22 (s, 2H), 4.03 (s, 2H), 3.02 (br s, 4H), 2.23 (s, 3H), 2.05-1.98 (m, 4H). **$^{13}$C NMR** (100 MHz, $CDCl_3$) δ 158.5, 139.8, 138.0, 137.9, 133.7, 130.1, 129.0, 128.9, 128.6, 127.9, 124.6, 116.5, 113.9, 69.3, 62.9, 58.9. 53.5, 50.8, 45.9, 23.1, 21.3. **HRMS (ESI$^+$)** calc. for $[C_{26}H_{29}NO_3S+H]^+$ 436.1941, obt. 436.1950.

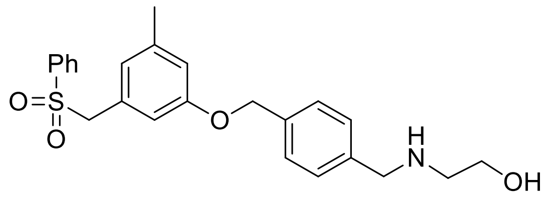


(**AMB-KA-036**)

Compound **KA-036 (Compound 36)** was prepared from **5a** and ethanolamine, purified using a 2-8% gradient of methanol in DCM, and obtained in 6% **yield** as a yellow oil. **$R_f$** (9:1 DCM:MeOH) ~0.59. **$^1$H NMR** (400 MHz, $CDCl_3$) δ 7.67 (dd, ill-defined, 2H), 7.65-760 (m, 1H), 7.56 (d, *J*=8.2 Hz, 2H), 7.50-7.43 (m, 4H), 6.75 (s, 1H), 6.54 (s, 1H), 6.48 (s, 1H), 4.97 (s, 2H), 4.24 (s, 2H), 3.81-3.71 (m, 2H), 3.06-3.01 (m, 1H), 2.96-2.91 (m, 1H), 2.23 (s, 3H). **$^{13}$C NMR** (100 MHz, $CDCl_3$) δ 158.5, 139.8, 138.0, 134.3, 133.7, 129.1, 128.9, 128.7, 127.9, 127.6, 124.7, 116.6, 113.9, 69.3, 62.9, 61.3, 54.1, 48.8, 21.3. **MS (ESI$^+$)** was unsuccessful.

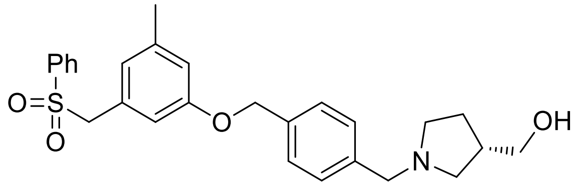


**AMB-KA-037**

Compound **KA-037 (Compound 37)** was prepared from **5a** and (*R*)-pyrrolidin-3-ylmethanol with acetic acid as a catalyst, purified using a methanol in DCM gradient over three columns, and obtained in 0.8% **yield**. **$R_f$** (9:1 DCM:MeOH) ~0.56. **$^1$H NMR** (400 MHz, $CDCl_3$): compound was too dilute for quantification from the observed peaks, and too dilute for a **$^{13}$C NMR** spectrum. **MS (ESI$^+$)** was unsuccessful.

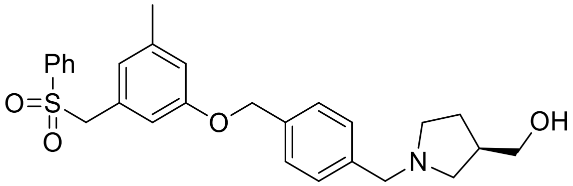


**AMB-KA-038**

Compound **KA-038 (Compound 38)** was prepared from **5a** and (*S*)-pyrrolidin-3-ylmethanol with acetic acid as a catalyst, purified using a 99:1 DCM:methanol eluent system, and obtained in 4% **yield**. $R_f$ (9:1 DCM:MeOH) ~0.42. **$^1$H NMR** (400 MHz, $CDCl_3$): compound was too dilute for quantification from the observed peaks, and too dilute for a **$^{13}$C NMR** spectrum. **MS (ESI$^+$)** was unsuccessful.

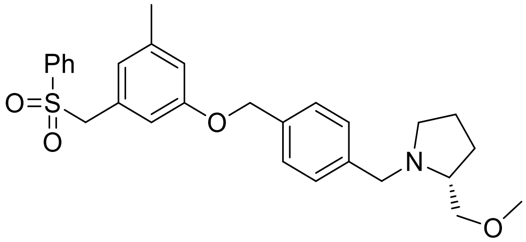


**AMB-KA-039**

Compound **KA-039 (Compound 39)** was prepared from *O*-methyl-D-prolinol with acetic acid as a catalyst, purified using a 99:1 DCM:methanol eluent system, and obtained in 21% **yield**. **$R_f$** (9:1 DCM:MeOH) ~0.79. **$^1$H NMR** (400 MHz, $CDCl_3$) δ 7.67 (dd, *J*=8.3 Hz, *J*=1.2 Hz, 2H), 7.62 (dd, *J*=7.4 Hz, *J*=7.4 Hz, 1H), 7.55 (d, *J*=8.0 Hz, 2H), 7.48 (dd, *J*=7.9 Hz, *J*=7.7 Hz, 2H), 7.40 (d, *J*=8.2 Hz, 2H), 6.75 (s, 1H), 6.52 (s, 1H), 6.49 (s, 1H), 5.60 (s, 1H), 4.95 (s, 2H), 4.24 (s, 2H), 3.55-3.47 (m, 2H), 3.41 (s, 2H), 3.23-3.15 (m, 1H), 2.70-2.65 (m, 1H), 2.58-2.51 (m, 1H), 2.23 (s, 3H), 2.04-1.95 (m, 1H), 1.81-1.67 (m, 2H), 1.57-1.48 (m, 1H). **$^{13}$C NMR** (100 MHz, $CDCl_3$) δ 158.6, 139.8, 138.0, 137.1, 134.6, 129.1, 128.8, 128.7, 127.8, 127.6, 127.4, 124.5, 116.7, 113.8, 69.4, 62.9, 60.5, 59.2, 58.7, 49.3, 27.8, 22.8, 21.3. **MS (ESI$^+$)** was unsuccessful.
